# ALFIN-LIKE Proteins Orchestrate H3K4me3-H3K27me3 Crosstalk to Regulate Plant Embryogenesis

**DOI:** 10.1101/2025.04.08.647773

**Authors:** Janik Scotton, Yixuan Fu, Edouard Tourdot, Marc W Schmid, Sara Simonini

## Abstract

In multicellular organisms such as animals and plants, development requires the precise regulation of gene expression, mediated not only by transcription factors but also by chromatin-based mechanisms. Among these, histone modifications like H3K4me3 and H3K27me3 play opposing roles in gene activation and repression, respectively. In *Arabidopsis thaliana*, H3K27me3 is deposited by the Polycomb Repressive Complex 2 (PRC2), while Trithorax group (TrxG) proteins mediate H3K4me3 deposition. While the functions of these writer complexes have been extensively studied, far less is known about the histone mark readers that interpret these modifications during development. Here, we investigate the antagonistic interplay between H3K27me3 and H3K4me3 during *Arabidopsis* embryogenesis. We identify a developmentally specific interaction between the FIS-PRC2 complex and ALFIN-LIKE (AL) proteins—a family of plant-specific PHD domain proteins that read H3K4me3. Our findings reveal a dynamic competition between these two marks during early embryogenesis that helps shape the epigenomic landscape of the developing seed. Disruption of AL function leads to severe developmental defects and loss of cell identity in early embryos. Moreover, loss of ALs impairs H3K4me3 deposition, resulting in aberrant spreading of H3K27me3, misregulation of developmental genes, and defects that persist into adult plant traits.

Together, our results show that proper embryonic development relies on a finely tuned antagonism between activating and repressive chromatin states—an interplay orchestrated not only by their writers but also by specific readers that translate these epigenetic cues into developmental outcomes.

## INTRODUCTION

Development relies on the precise regulation of gene expression to ensure that undifferentiated cells acquire and maintain specialized identities. This regulation extends beyond the DNA sequence to include chromatin-based mechanisms. Among these, histone modifications play a pivotal role in defining transcriptional states by modulating chromatin compaction and accessibility (Millán-Zambrano et al., 2022). These modifications are deposited by specialized protein complexes known as "writers," which enzymatically modify specific amino acid residues within histone tails. Two of the most extensively studied histone modifications are the methylation of lysine 4 and lysine 27 in the tail of histone 3 tail, referred to as H3K4me3 and H3K27me3, respectively. These modifications are antagonistic and non mutually exclusive, with H3K27me3 linked to gene repression and H3K4me3 to activation. The *Polycomb* Repressive Complex 2 (PRC2; Cao et al., 2002) is the writer for H3K27me3 and catalyzes mono-, di-, and trimethylation at this residue to establish repressive chromatin domains. PRC2 composition and function are highly conserved among multicellular organisms, from *Drosophila* to humans and plants (Grossniklaus and Paro, 2014). Its function is essential for development, as it ensures the correct gene expression patterns of developmental and cell identity genes. In flowering plants, the genes encoding PRC2 subunits have undergone duplication. In the model plant *Arabidopsis thaliana*, three methyltransferases - MEDEA (MEA), CURLYLEAF (CLF), and SWINGER (SWN) - assemble into at least three specialized PRC2 complexes, each with distinct core and accessory subunits (Grossniklaus and Paro, 2014). The MEA-PRC2 (also called FIS-PRC2) complex functions specifically in the female gametophyte, early embryo, and endosperm, a placental-like tissue that fills the seed cavity and provides nutrients to the growing embryo (Chaudhury et al., 1997; Grossniklaus et al., 1998; Köhler et al., 2003; Kiyosue et al., 1999; Ohad et al., 1996). By contrast, CLF- and SWN-based PRC2 complexes regulate post-embryonic transitions, specifically the trasnition from embryo to seedling transition and the vernalization response (Bastow et al., 2004; Bouyer et al., 2011; Schubert et al., 2006). The importance of PRC2 in development is particularly evident during embryogenesis and stage transitions, where cells must make critical developmental decisions. In *Arabidopsis*, loss of PRC2 function leads to severe developmental defects, such as embryo abortion in *mea* mutants (Grossniklaus et al., 1998; Simonini et al., 2021). Simultaneous *clf* and *swn* mutations trigger immediate tissue dedifferentiation after germination (Chanvivattana et al., 2004).

Trithorax group (*TrxG*) proteins are the "writers" of H3K4me3, a histone modification essential for gene expression. It regulates cell fate, development, and embryogenesis, with misregulation is linked to tumorigenesis in animals (Ingham, 1983; Schuettengruber et al., 2011). The functions of *TrxG* genes in *Arabidopsis* and other plant models remain poorly understood. Although mutations in the Arabidopsis *TrxG* homolog ATX1 do not affect embryogenesis (Alvarez-Venegas et al., 2003; Pien et al., 2008), loss of the TrxG protein *TRAUCO* disrupts H3K4me3 deposition and leads to embryo abortion (Aquea et al., 2010). This suggests that proper H3K4me3 deposition is critical for embryonic growth and patterning, though additional evidence is scarce and the underlying mechanisms remain unclear.

In addition to methylation, other histone post-translational modifications, such as acetylation and phosphorylation, also regulate gene expression. Unlike these modifications, which alter the electronic charge of the histone tail and affect its chemical properties, methylation does not change the substrate’s charge. As a result, histone lysine methylation is primarily thought to function by recruiting specific reader and effector proteins that recognize distinct methylated sites (Berger, 2007; Millán-Zambrano et al., 2022). For instance, H3K4me3 promotes transcription by recruiting ATP-dependent chromatin remodelers, which enhance chromatin accessibility by repositioning or evicting nucleosomes thus facilitating access to the transcriptional machinery to promoter regions (Bayona-Feliu et al., 2021; Li et al., 2015; Tosi et al., 2013).

To understand how histone methylation translates into specific developmental outcomes, it is therefore essential to identify reader proteins and determine their specific roles. Although histone methylation writer complexes have been extensively characterized in animals, their characterization in plants remains less developed, and identifying the functional readers of histone modification remains an even greater challenge. Unlike writers, which are typically encoded by a small number of genes, reader proteins often belong to large multigene families, sometimes with dozens of members (Hyun et al., 2017; Yun et al., 2011). This complexity makes it difficult to determine their precise developmental roles, complicating functional studies. Among these families, plant homeodomain (PHD) fingers are among the best characterized in *Arabidopsis* (De Lucia et al., 2008; Gao et al., 2023; Kim and Sung, 2017; Lai et al., 2020; Molitor et al., 2014; Mouriz et al., 2015; Sung et al., 2006). While their biochemical binding properties are well understood, their developmental significance remains unclear, as functional redundancy and compensatory mechanisms may mask loss-of-function phenotypes. Moreover, their influence on global histone methylation dynamics during development is not well understood.

In this study, we investigate the functional relationship between H3K27me3 and H3K4me3 during *Arabidopsis* embryogenesis and early seed development. We identify a developmental-stage-specific interaction between FIS-PRC2 subunits and the H3K4me3 readers ALFIN-LIKE (ALs), a class of plant-specific PHD domain proteins that bind H3K4me3 and whose developmental role is only beginning to emerge (Jin et al., 2024; Kayum et al., 2015; Molitor et al., 2014; Peng et al., 2018; Song et al., 2013; Su et al., 2025; Wei et al., 2015; Xu et al., 2024). Our findings reveal that during early embryogenesis, H3K27me3 and H3K4me3 engage in a dynamic ‘tug-of-war,’ competing for genomic occupancy. This extensive activity helps establish the epigenomic landscape in the early embryo and contributes to proper development. Disrupting H3K4me3 occupancy leads to the spreading of H3K27me3 into those regions that lose H3K4me3, which affect embryonic gene expression and plant development. Our results highlight that proper development does not simply rely on the individual functions of epigenetic machineries, but rather on their coordinated, antagonistically interplay, ensuring that neither dominates and that gene expression remains balanced.

## RESULTS

### PRC2 interactome analysis reveals a potential crosstalk with the antagonistic histone mark H3K4me3

The PRC2 plays a central role in establishing and maintaining gene repression through H3K27me3 deposition. However, because PRC2 lacks intrinsic DNA-binding activity, the mechanisms guiding its recruitment to target loci remain poorly understood, particularly in plants. To identify potential PRC2 interactors that may facilitate recruitment or provide insight into the chromatin environment in which PRC2 operates, we performed immunoprecipitation with liquid chromatography and tandem mass spectrometry (IP LC-MS/MS; Fig. 1A). We used *Arabidopsis* PRC2 methyltransferases MEA-GFP (Simonini et al., 2021), CLF-GFP, (Shu et al., 2019) and SWN-GFP (Shu et al., 2019), each expressed under its endogenous promoter in their respective mutant backgrounds, as bait. Given their distinct roles, we performed this analysis on 14-day-old seedlings for CLF and SWN, while for MEA we analyzed flowers 1-2 days post anthesis.

**Figure 1.**
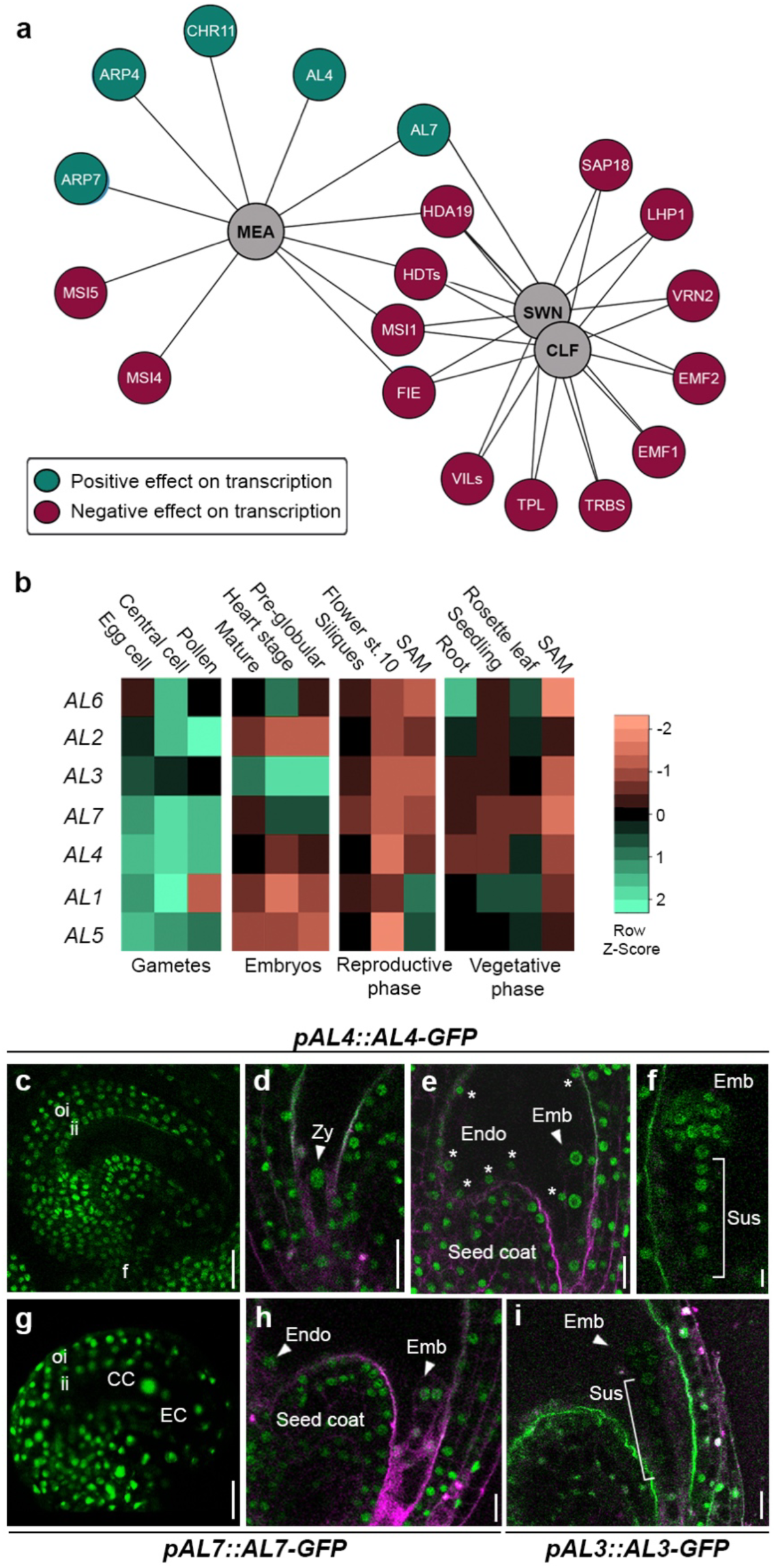
Members of the ALFIN-LIKE protein family immunoprecipitate with PRC2. **(a)** STRING network representing the interactome of MEA, CLF, and SWN based on IP/MS-MS analysis. **(b)** Expression dynamics of *AL* genes across different plant tissues. **(c-f)** Confocal images of *pAL4::AL4-GFP* expression in a mature ovule (c), the zygote (d), developing embryos (e-f), and the endosperm (e). **(g-h)** Confocal images of *pAL7::AL7-GFP* expression in a mature ovule (g), showing strong signal in the female gametes (central cell and egg cell), and in the developing embryo and endosperm (h). **(i)** Confocal image of *pAL3::AL3-GFP* expression in the developing embryo. EC: egg cell; CC: central cell; Emb: embryo; End: endosperm; Sus: suspensor; ii: inner integument; oi: outer integument; f: funiculus. Scale bar: 20 µm.

As expected, our analysis identified the core PRC2 subunits FERTILIZATION INDEPENDENT ENDOSPERM (FIE, (Ohad et al., 1996)) and MULTICOPY SUPPRESSOR OF IRA1 (MSI1, (Köhler et al., 2003)) in all three immunoprecipitates, along with complex-specific factors such as REDUCED VERNALIZATION 2 (VRN2, (Gendall et al., 2001)) and EMBRYONIC FLOWER 2 (EMF2, (Yoshida et al., 2001)), reinforcing the distinct PRC2 variants (Fig. 1a, Supplementary Table 1). The strong overlap between CLF- and SWN-associated proteins, including VIN3-LIKE1/2 (VIL1/2), TOPLESS (TPL, (Baile et al., 2020; Long et al., 2006)), and TELOMERE REPEAT BINDING FACTORS (TRBs, (Zhou et al., 2018)), supports their functional redundancy during development. Intriguingly, MEA strongly associated with MSI4 and MSI5 rather than MSI1, suggesting a specific composition of MEA-PRC2 (Fig. 1A). All three PRC2 methyltransferases also co-purified with histone deacetylases such as HISTONE DEACETYLASE19 (HDA19, (Tian et al., 2003; Wu et al., 2000)) and members of the HISTONE DEACETYLASE (HDT) family (Sridha and Wu, 2006), which collaborate with PRC2 to maintain gene repression through histone deacetylation and H3K27me3 deposition (Godwin and Farrona, 2022).

Unexpectedly, MEA-PRC2 also co-purified with components of chromatin remodeling complexes typically associated with gene activation, including ACTIN-RELATED PROTEIN 4/7 (ARP4/7, (Meagher et al., 2010)) and CHROMATIN-REMODELING PROTEIN 11 (CHR11, (Huanca-Mamani et al., 2005)). Strikingly, MEA also interacted with two ALFIN-LIKE (AL) proteins, AL4 and AL7, which are known H3K4me3 readers (Fig. 1a, Supplementary Table 1, (Liang et al., 2018; Su et al., 2025)). SWN-GFP also co-purified with AL7, but not AL4. This suggests a potential crosstalk between H3K4me3 and H3K27me3, which appears to be more pronounced in MEA-PRC2, raising the possibility that this interaction is developmentally regulated and particularly relevant during early seed development, where MEA is active. These findings also hint at distinct chromatin environments for MEA-PRC2 versus CLF/SWN-PRC2.

To determine whether AL proteins could functionally interact with MEA-PRC2, we examined their expression domains. Publicly available transcriptome data confirmed that *AL* genes, including *AL4* and *AL7*, are expressed in the embryo and endosperm (Fig. 1b), tissues where MEA is expressed. More broadly, ALs proteins are widely expressed across various plant tissues and developmental stages (Fig. 1b). To validate their spatial expression, we generated GFP-tagged reporters for AL3, AL4, and AL7 under their endogenous promoters.

Consistent with transcriptomic data, AL-GFP fusions showed fluorescence in the ovule and in the female gametes central cell and egg cell (Fig. 1c-i). After fertilization, ALs were detected in the zygote, embryo, and suspensor throughout embryogenesis (Fig. 1d-i), confirming their presence in the same cell types where MEA-PRC2 operates ((Grossniklaus et al., 1998; Kiyosue et al., 1999; Vielle-Calzada et al., 1999). These findings point to a previously unrecognized potential relationship between PRC2 and factors associated with the antagonistic H3K4me3 mark. While PRC2 is classically viewed as a repressive complex, our data might suggest that at least in the MEA context, PRC2 operates within a chromatin landscape enriched in both activating and repressive marks, hinting at a dynamic regulatory mechanism occurring during seed development.

### ALs perform essential function during reproduction

The *Arabidopsis* genome encodes seven AL proteins (AL1–AL7), which bind H3K4me3 in vitro and in vivo and have emerged as key component of chromatin remodeling complexes (Su et al., 2025). AL proteins have been implicated in ABA signaling, seed germination, and stress responses (Jin et al., 2024; Jing et al., 2025; Kayum et al., 2015; Wei et al., 2015), however, their broader developmental significance remains unclear due to challenges in generating higher-order mutants. Septuple al mutant have been shown to develop with extremely severe pleiotropic defects (Su et al., 2025; Xu et al., 2024), emphasizing their importance but also complicating functional dissection and the understanding of their function in specific developmental pathways. To investigate the biological and developmental function of ALs proteins, we first generated the *al4 al7* double mutant, as these two family members co-purified with MEA in our IP LC-MS/MS analysis. Since a T-DNA insertional mutant was available for *AL7* but not *AL4*, we used CRISPR-Cas9 to introduce mutations in *AL4* within the *al7* mutant background. However, the *al4 al7* double mutant displayed no obvious phenotypic differences compared to wild-type plants in any examined tissues. This result, while unsurprising, suggests functional redundancy among the seven AL genes, consistent with their overlapping expression patterns and putative similar roles.

To overcome this redundancy, we aimed to generate a septuple *al* mutant (*al1/2/3/4/5/6/7*) using a multi-guide RNA CRISPR-Cas9 system to simultaneously target *AL1*, *AL2*, *AL3*, *AL5*, and *AL6* in the *al4 al7* background (Fig. EV1a). Our initial screening retrieved deleterious mutations in *AL1*, *AL3*, *AL5*, and *AL6*, but not in *AL2*. The absence of *AL2* mutants was likely due to guide RNA inefficiency, as we failed to detect any *AL2* deleterious or in-frame mutations across hundreds of indipendent transformants. To address this, we transformed the *al3/4/5/6/7* quintuple mutant with a new *AL2* guide RNA, successfully obtaining the *al2/3/4/5/6/7* sextuple mutant. However, we failed to recover a full septuple mutant (*al1/2/3/4/5/6/7*) or even a sextuple mutant carrying *AL2* in heterozygosity. The inability to generate a complete *al1/2/3/4/5/6/7* mutant strongly suggests that total loss of AL function is lethal, a conclusion supported by two recent studies reporting similar findings using high-order CRISPR-Cas9-generated al mutants ((Su et al., 2025; Xu et al., 2024).

Given the different timings at which we obtained the two sextuple mutant backgrounds (*al1/3/4/5/6/7* and *al2/3/4/5/6/7*), the majority of our analyses focused on *al1/3/4/5/6/7*, the first successfully generated sextuple mutant. Importantly, the later obtained *al2/3/4/5/6/7* displayed no distinct phenotypes compared to *al1/3/4/5/6/7*, confirming that our findings reflect the function of the AL protein family as a whole.

To dissect the developmental impact of AL loss, we systematically analyzed plants carrying triple, quadruple, quintuple, and sextuple mutations. Triple (*al4/5/7*) and quadruple (*al4/5/6/7*, *al3/4/5/7*) mutants largely resembled wild-type plants (Figs. 2a and EV1b). The first mild developmental defects emerged in the *al1/4/5/6/7* and *al3/4/5/6/7* quintuple mutants, which became significantly exacerbated in *al1/3/4/5/6/7* sextuple mutants (Figs. 2a and EV1b). At a macroscopic level, quintuple and sextuple mutants exhibited altered flowering time, slow development, reduced plant height, fewer branches, longer internodes, and fewer flowers with disrupted phyllotaxis (Figs. 2a and EV1b). Additionally, siliques were shorter, suggesting fertilization defects (Figs. 2a and EV1b-c). These phenotypes were also present in *al1/2/3/4/5/7* plants (Fig. EV1d).

**Figure 2.**
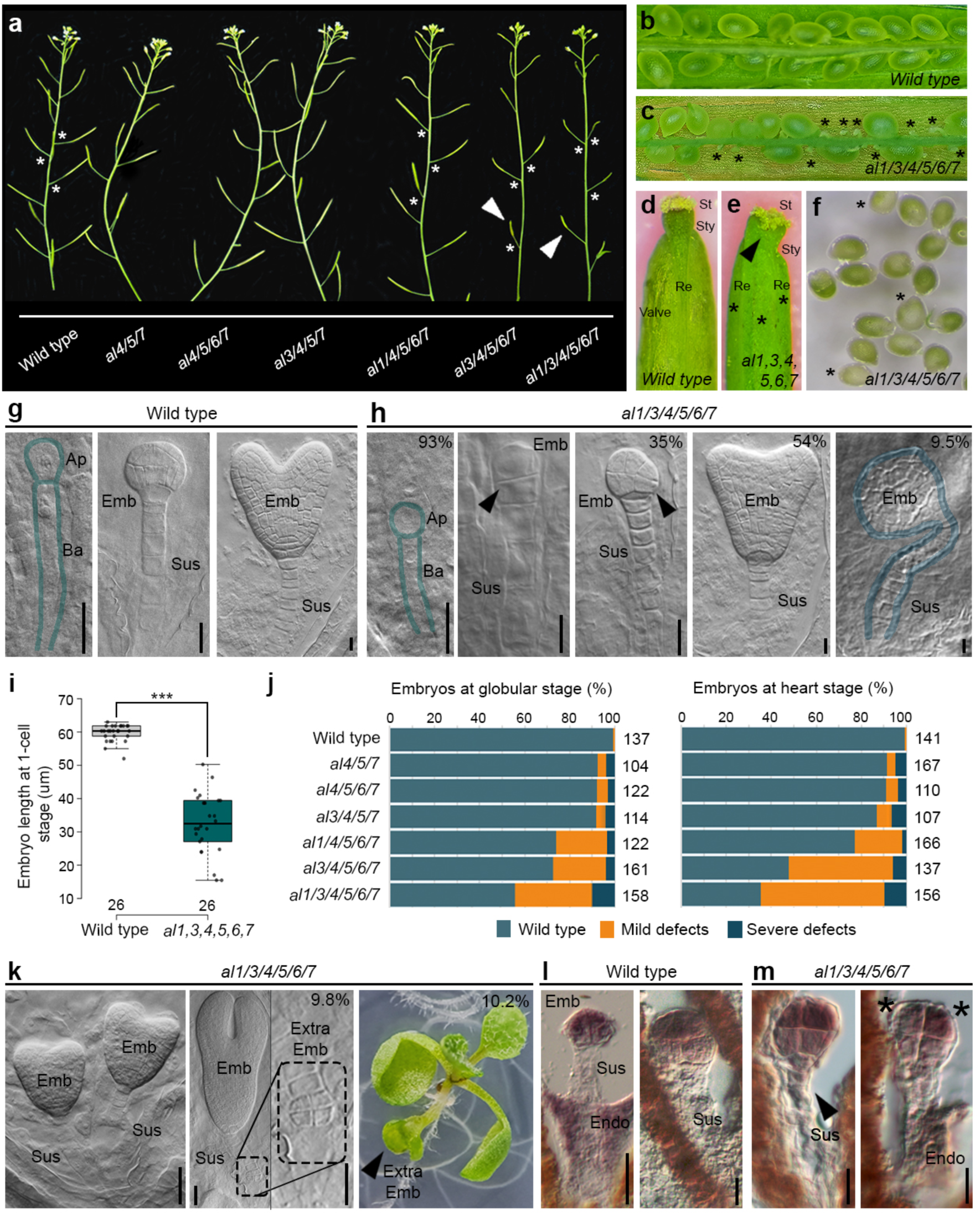
ALs are required for proper embryogenesis. **(a)** Primary stems of wild-type and *al* triple, quadruple, quintuple, and sextuple mutants, showing reduced flower number, altered phyllotaxis (asterisks), and shorter siliques (arrow head). **(b-c)** Close-up of open siliques from wild-type (B) and *al* sextuple mutant (C), highlighting ovule abortion (asterisks). **(d)** Isolated seeds from the *al* sextuple mutant, showing visible embryos inside most seeds, except in a few where embryonic growth is arrested and the seed is white (asterisks). **(e-f)** Close-up of carpel tips from wild-type (E) and *al* sextuple mutant (F), showing an increased number of valves (asterisks) and a cleft in the style (arrow head). **(g-h)** Clearing images of wild-type (G) and alfin sextuple mutant (H) embryos at different developmental stages. The percentage of embryos displaying these defects is indicated in the top right corner. **(i)** Quantification of embryo elongation at the one-cell stage in wild-type and *al* sextuple mutants. **(j)** Quantification of patterning defects in % of embryo showing a given phenotype in wild type (J) and *al* sextuple mutant (K) embryos at the globular and heart stages based on the clearing analysis. **(k)** Ectopic embryo developmenrt in *al* sextuple mutants, during different developmental stages. The percentage of these embryos among the *al* sextuple offspring is indicated in the top right corner. **(l-m)** *In situ* hybridization using a *WOX2* antisense probe in wild-type (L) and *al* sextuple mutant (M) embryos. Arrow heads indicate ectopic exprssion of *WOX2* in the suspensor of *al* sextuple mutant embryos. Asterisks indicartes two cells in the upper tier of the *al* sextuple embryo that express different levels of *WOX2*. Ap: apical cell; Ba: basal cell; St: stigma; Sty: style; Re: replum; Emb: embryo; Sus: suspensor; Extra Emb: extraembryonic embryo. Scale bar: 20 µm.

Given our focus on MEA-PRC2 function, we examined the *al1/3/4/5/6/7* sextuple mutant in more detail to assess potential seed-related defects. Siliques from this mutant contained numerous aborted ovules (Fig. 2b-c, suggesting impaired fertilization, as well as a subset of white seeds, indicative of embryo abortion (Fig. 2d).

A possible cause of the first defect - ovule abortion - was linked to abnormal pistil morphology, with sextuple mutants developing four valves instead of two and displaying an elongated cleft in the style (Fig. 2d-e). Given the role of the style in guiding pollen tube growth, these structural defects likely contributed to fertilization failure. To further assess reproductive defects, we performed manual pollination using both wild-type and self-mutant pollen on *al1/3/4/5/6/7* carpels. While this partially rescued seed set, it failed to restore wild-type levels, suggesting additional defects in female gametogenesis (Fig. EV2a).

### AL proteins regulate early embryonic cell fate

Post-fertilization, we observed the most striking phenotypes in *al* multiple mutants. In wild-type, the zygote elongates before undergoing its first asymmetric mitotic division (Fig. 2g), generating a small apical cell (giving rise to the embryo proper) and a larger basal cell (which forms later the suspensor, an extra-embryonic structure connecting the embryo to maternal tissues) (ten Hove et al., 2015). In contrast, *al1/3/4/5/6/7* mutant embryos failed to elongate as in wild type and underwent a more symmetric first division (Fig. 2h-i). Later, aberrant division planes in the embryo proper resulted in a spectrum of developmental abnormalities, ranging from mild defects to severe embryo abortion (Figs. 2h, j and EV2b). In rare cases, we observed twin embryos, each with a well-patterned structure and its own suspensor (Figs. 2k and EV2c). These twins could arise via two mechanisms: (i)- Division of the zygote occurring post-fertilization, where a single fertilized egg splits into two embryos, or (ii)- The presence of two egg cells in a single female gametophyte, where each sperm fertilizes a separate egg, leaving the central cell unfertilized. Supporting the first hypothesis, the detected twin embryos often co-developed with a functional endosperm (Fig. 2k). However, we also observed seeds with twin embryos lacking a functional endosperm and ovules harboring two egg cells (Fig. EV2c-d), suggesting that both mechanisms may contribute to this phenotype.

Strikingly, patterning defects extended beyond the embryo proper to the suspensor, which frequently displayed excessive cell divisions, occasionally giving rise to an additional embryo (Figs. 2h, k and EV2e). These suspensor-derived embryos varied in morphology, ranging from aberrant clusters to well-patterned structures that developed into mature embryos. However, these embryos were generally delayed in development and frequently fused at the hypocotyl level to the main seedling (Fig. 2k). Additionally, we also detected mild embryonic defects including the formation of three cotyledons instead of two (Fig. EV2f). All these embryonic phenotypes were also observed in sextuple mutant combination *al1/2/3/4/5/7* and - albeit less frequently - in quadruple and quintuple mutants (Figs. 2j and EV2a-b,g).

Our morphological analyses indicate that AL proteins are essential for proper cell identity acquisition in both the apical and basal cells immediately after fertilization. In their absence, the basal cell fails to specify suspensor identity, instead retaining embryonic potential and forming ectopic embryos. This conclusion is further supported by the misexpression of the embryo-specific gene *WOX2* (Haecker et al., 2004) (Fig. 2l-m). In wild-type embryos, *WOX2* expression is strictly confined to the embryo proper, predominantly in the upper tier (Fig. 2l). However, in *al1/3/4/5/6/7* mutants, *WOX2* is ectopically expressed in the suspensor (Fig. 2m), confirming that suspensor cells acquire embryonic identity. Additionally, *WOX2* expression in the upper tier was irregular, with individual cells displaying varying expression levels, further suggesting disrupted cell identity specification.

Altogether, these findings establish AL proteins as key regulators of early embryogenesis, particularly in apical-basal cell fate specification. Given that ALs are H3K4me3 readers, these observations also suggest that H3K4me3 presence in the zygote and early embryo - and its correct interpretation - are critical for cell fate specification during early embryogenesis.

### ALs have tissue-specific interactomes

Recent studies identified AL proteins as partners of chromatin remodelers and transcriptional activators (Su et al., 2025), but this was based on seedling tissue. Since the strongest AL-related phenotypes occur during seed development, we investigated whether AL proteins have tissue-specific interactomes and stage-dependent proximity to PRC2. To explore this, we performed immunoprecipitation followed by mass spectrometry (IP/MS) using AL4-GFP in young siliques at 1–4 days after fertilization. We introduced AL4-GFP into the *al1/3/4/5/6/7* sextuple mutant background, where it successfully complemented developmental defects, restoring phenotypes similar to the *al1/3/5/6/7* quintuple mutant (Fig. EV3A). This confirmed the functionality of the AL4-GFP fusion protein. Additionally, performing IP/MS in the absence of five other AL family members helped identify ALs-specific interactors, reducing redundancy that often complicates multi-protein family analyses.

Our IP/MS analysis revealed a strong enrichment of chromatin-associated factors (Fig. 3 and Supplementary Table 1), consistent with previously reported AL function. Specifically, AL4 immunoprecipitates contained subunits of nucleosome remodeling complexes, including BRAHMA (BRM, (Farrona et al., 2011)), CHR4, CHR12, CHR17, CHR23, and components of the SWI complex (Supplementary Table 1). Additionally, ARP4 and ARP7 - previously detected in MEA-GFP immunoprecipitates - were also present, reinforcing the functional overlap between these complexes. Beyond factors associated with gene activation, we identified several repressors, including the PRC2 subunit MSI1, the PRC2 recruiters VAL1, VAL3, and BPC6, as well as histone deacetylases and PRC1 subunits such as RING1A. Notably, PRC1 was the first identified AL interactor, further supporting the idea that AL proteins operate within a dynamic chromatin environment where epigenetic states are finely regulated. Many of these interactors were also found in AL4 immunoprecipitates from seedlings. However, a distinctive feature of AL4 interactomes in developing seeds was the enrichment of DNA replication and repair factors, including components of the MINICHROSOME MAINTENANCE (MCM) family (Fig. 3 and Supplementary Table 1). This suggests that AL proteins may function in proximity to the replication fork, potentially engaging with modified histones during DNA replication or ensuring faithful epigenetic inheritance across cell divisions.

**Figure 3.**
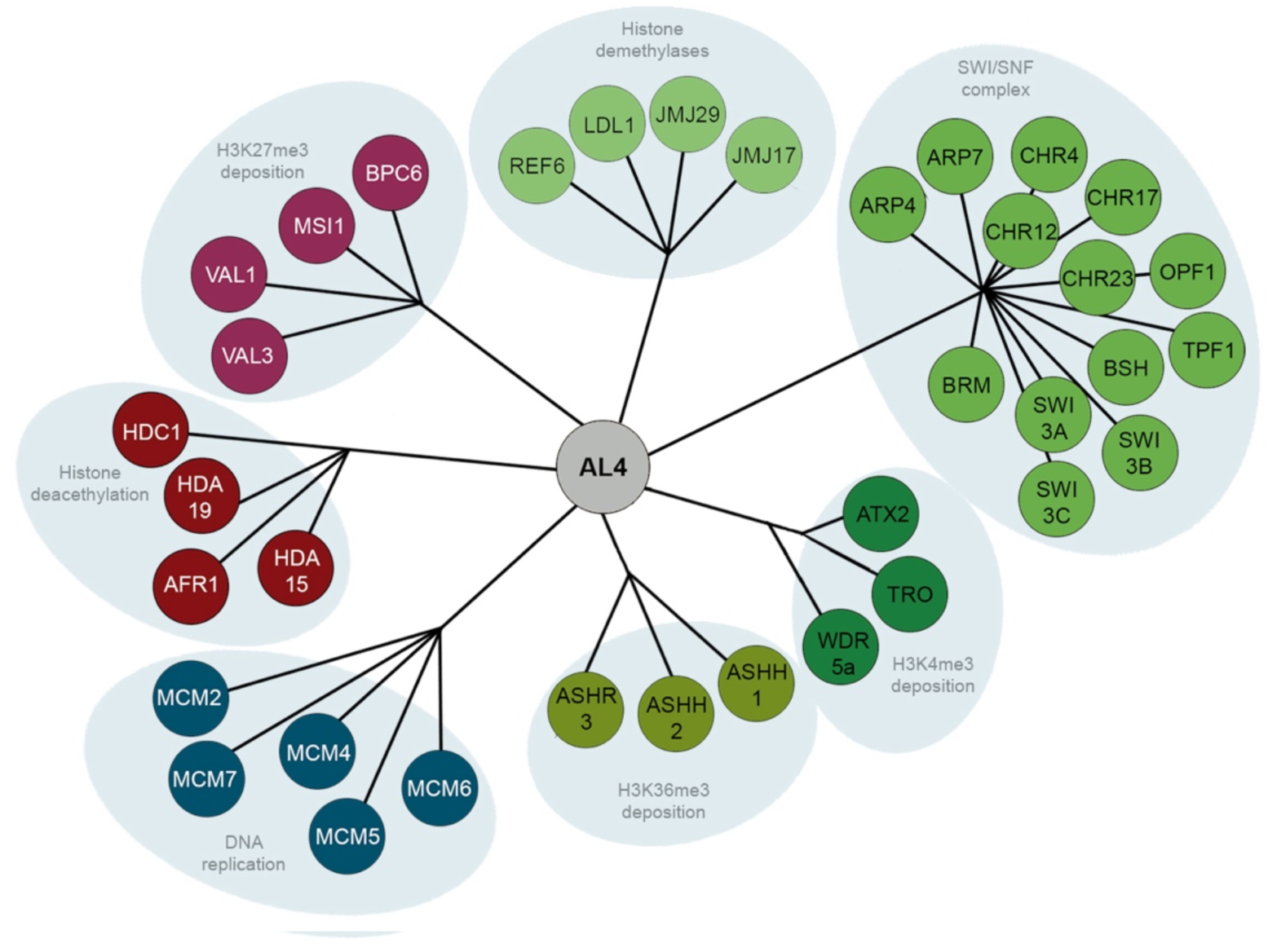
Interactome of AL4 in dveloping seeds. Graphic representation of the AL4 interactome based on our IP/MS experiment, highlighting key interactors and their molecular functions.

Altogether our IP/MS analysis provides further evidence for a potential interplay between establishment and maintenance of H3K4me3 and H3K27me3 domains during the early stages of seed development.

### ALs are required for correct expression of developmental genes

To investigate the impact of ALs function on transcription and development, we performed RNA-Seq transcriptome analysis on developing seeds from 1 to 4 days after fertilization, thus covering embryonic stages from the zygote to early globular stage. These early stages mark the onset of tissue specification, where pluripotent cells gradually commit to distinct developmental trajectories. We compared wild-type seeds with the *al3/4/5/6/7* quintuple mutant and the *al1/3/4/5/6/7* sextuple mutant, as embryonic defects are already evident in the quintuple mutant but become more severe in the sextuple mutant. This comparative approach increased the confidence of our findings and allowed us to better define the genetic pathways involved in early embryogenesis.

Differentially expressed genes (DEGs) were categorized as upregulated or downregulated (Fig. 4a, Supplementary Table 2), and further grouped by clustering analysis allowing the identification of distinct expression patterns (Fig. 4b). Among downregulated genes, some showed a progressive decrease in expression from the quintuple to the sextuple mutant, while others were almost completely silenced in both backgrounds. Given that H3K4me3 and AL proteins are generally associated with transcriptional activation, we propose that core AL target genes are predominantly among the downregulated DEGs. For upregulated genes, clustering analysis identified three distinct patterns: (i) genes exclusively upregulated in the sextuple mutant, (ii) genes that progressively increased from the quintuple to the sextuple mutant, and (iii) genes stably upregulated in both backgrounds (Fig. 4c and Supplementary Table 2). Gene Ontology (GO) enrichment analysis of downregulated genes highlighted biological processes related to embryo and seed development, as well as cell identity acquisition (Fig. 4c and Supplementary Table 2), consistent with the observed mutant phenotypes. In contrast, GO analysis of upregulated genes identified distinct biological processes, including hormonal responses—particularly auxin signaling—and developmental processes beyond seed formation, such as flower development and reproductive functions like pollen tube growth (Fig. 4d and Supplementary Table 2).

**Figure 4.**
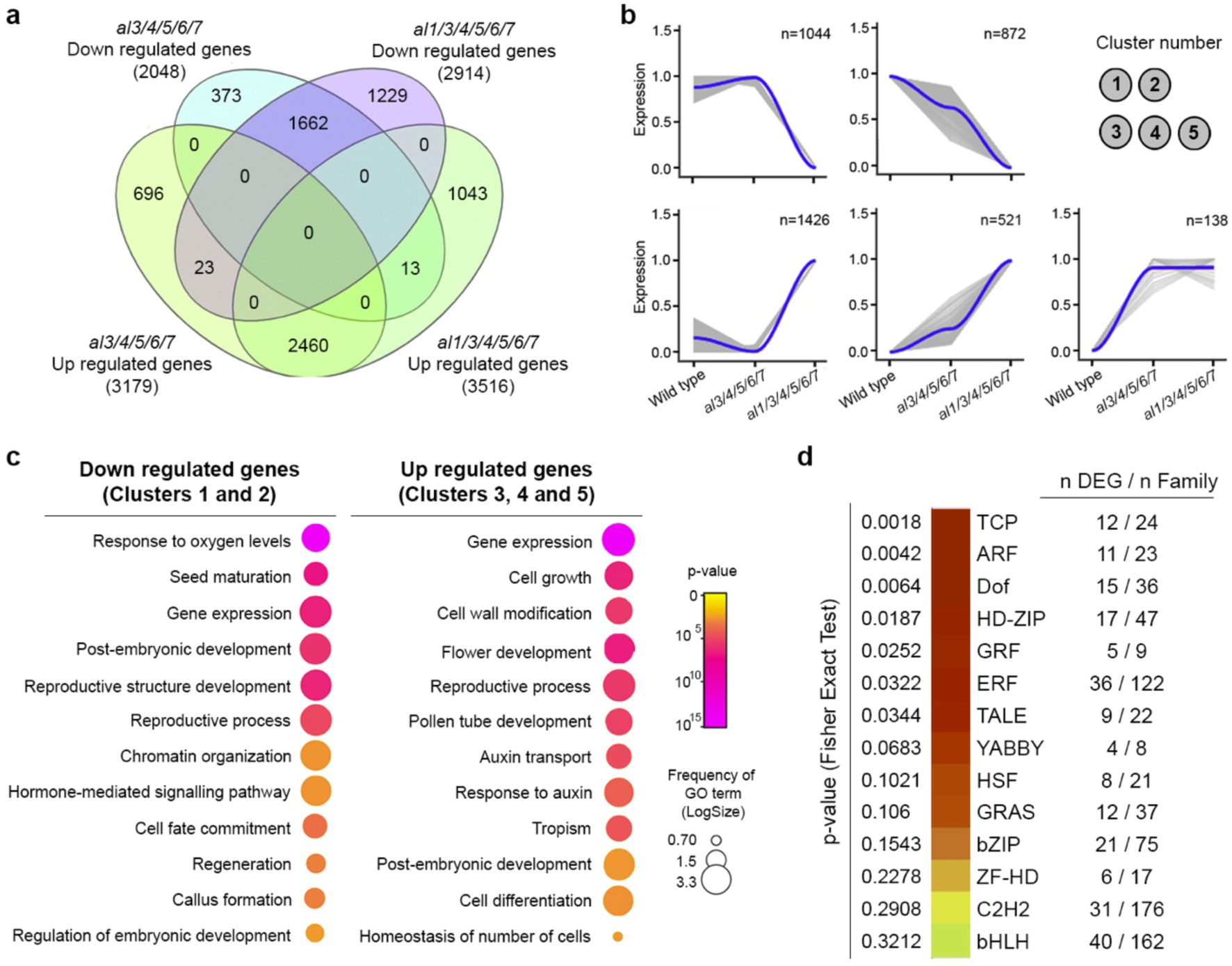
Transcriptome analysis of *al* multiple mutants reveals a close link with developmental processes and transcriptional regulation. **(a)** Venn diagram depicting the overlap in RNA-Seq analysis of wild-type, *al3/4/5/6/7*, and *al1/3/4/5/6/7* young seeds. **(b)** Cluster analysis showing five clusters with distinct expression patterns, with downregulated genes in clusters 1 and 2 and upregulated genes in clusters 3, 4, and 5. **(c)** GO term enrichment analysis of upregulated genes (clusters 3, 4, and 5) and downregulated genes (clusters 1 and 2). The color of each dot corresponds to the p-value, and the size of the circle represents the frequency of that particular GO term, shown as LogSize. **(d)** Enrichment of transcription factor families with deregulated expression in *al1/3/4/5/6/7* seeds. The p-value was calculated using Fisher’s exact test. The number of deregulated genes within each family is indicated in the top right corner.

These findings reinforce the role of AL proteins as key regulators of developmental programs, particularly during embryo development. We hypothesize that upregulated genes reflect secondary effects arising from the deregulation of primary AL targets.

Interestingly, the term “regulation of gene expression” emerged as a highly represented and shared GO term among both upregulated and downregulated genes, suggesting that AL proteins function as central regulators of transcriptional cascades, likely influencing the activity of master transcription factors (TFs). Indeed, several TF families were significantly enriched among DEGs, including TCPs, ARFs, HD-ZIP, GRF and TALE – all key regulators of plant development and morphogenesis (Fig. 4d and Supplementary Table 2). Altoghether, these results highlight the critical role of AL proteins in orchestrating early embryogenesis by regulating key transcriptional networks.

### AL proteins regulate histone methylation dynamics during early seed development

AL proteins recognize H3K4me3 marks, and their loss would be expected to impact gene expression while only mildly affecting H3K4me3 deposition. However, immunoprecipitation of AL4-GFP revealed interactions with H3K4me3 writer complexes, suggesting a potential role in maintaining epigenetic memory across cell divisions. To investigate this, we performed CUT&Tag on isolated seeds (1-4 days after fertilization) to map H3K4me3 occupancy in wild-type, *al3/4/5/6/7* quintuple, and *al1/3/4/5/6/7* sextuple mutants (Fig. 5a-b and Supplementary Table 3). The results showed that AL loss disrupted H3K4me3 deposition, leading to both reduced and ectopically gained marks at specific genomic regions (Fig. 5a-b and Supplementary Table 3). This suggests that AL proteins not only recognize H3K4me3 to activate transcription but also help maintain epigenetic states during development.

**Figure 5.**
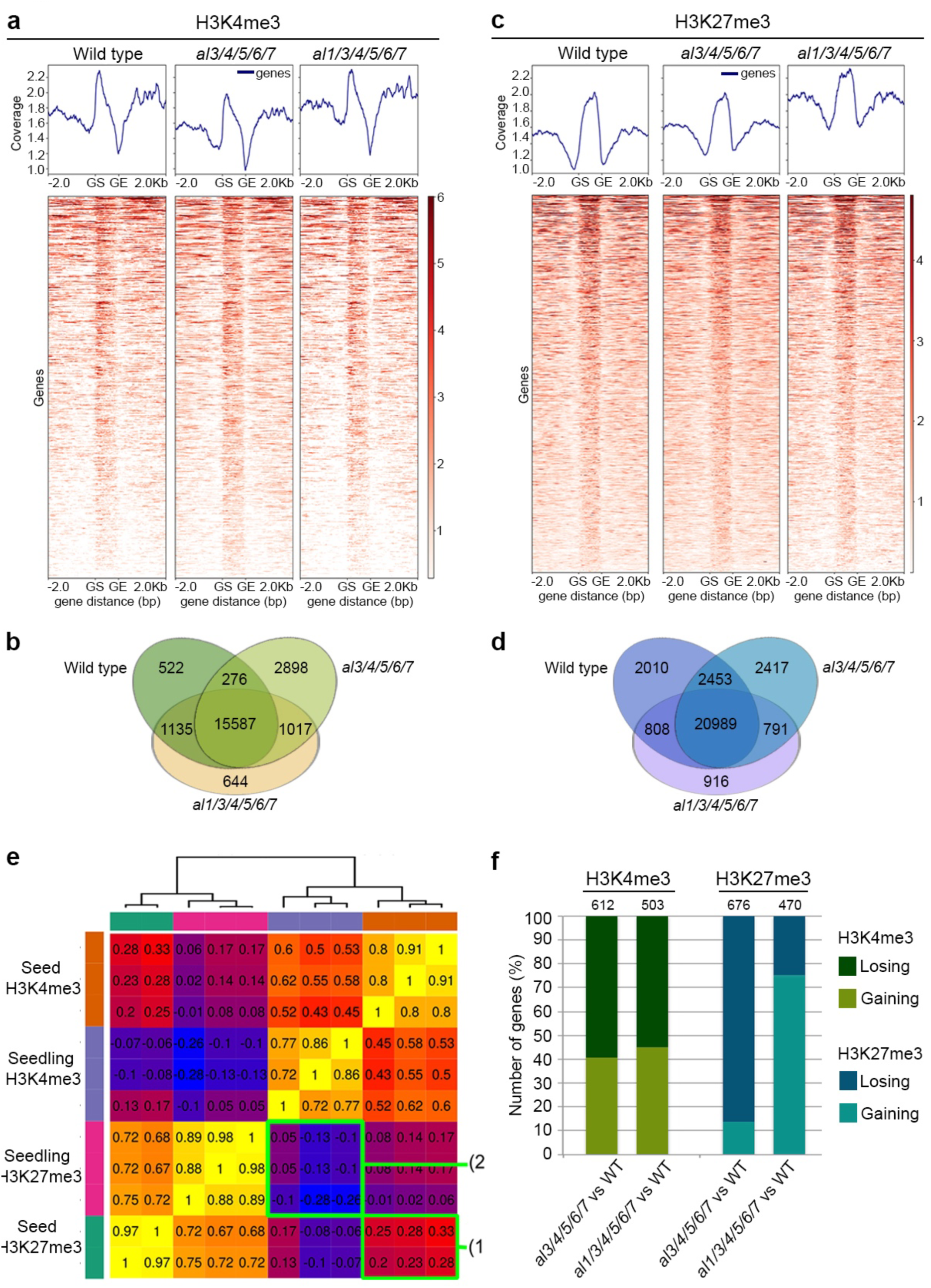
AL proteins coordinate H3K4me3 and H3K27me3 occupancy. **(a)** Heatmap and composite profiles showing H3K4me3 occupancy around gene loci in wild-type, *al3/4/5/6/7*, and *al1/3/4/5/6/7* young seeds. GS: gene start; GE: gene end. **(b)** Venn diagram summarizing H3K4me3 occupancy identified by CUT&Tag in wild-type, *al3/4/5/6/7*, and *al1/3/4/5/6/7* young seeds. **(c)** Heatmap and composite profiles showing H3K27me3 occupancy around gene loci in wild-type, *al3/4/5/6/7*, and *al1/3/4/5/6/7* young seeds. GS: gene start; GE: gene end. **(d)** Venn diagram summarizing H3K27me3 occupancy identified by CUT&Tag in wild-type, *al3/4/5/6/7*, and *al1/3/4/5/6/7* young seeds. **(e)** Heatmap showing sample correlation among CUT&Tag datasets for H3K4me3 and H3K27me3 in wild-type seedlings and seeds. Square 1: correlation between H3K4me3 and H3K27me3 in wild-type seeds. Square 2: correlation between H3K4me3 and H3K27me3 in wild-type seedlings. Higher correlation in seeds suggests greater similarity between the two histone marks compared to seedlings. **(f)** Bar chart showing the percentage of genes gaining or losing H3K4me3 and H3K27me3 in *al3/4/5/6/7* and *al1/3/4/5/6/7* young seeds relative to wild type.

Given the potential interplay between AL proteins and PRC2, we also profiled H3K27me3 occupancy via CUT&Tag (Fig. 5c-d), while considering two possible models: (i) ALs recruit PRC2 to specific loci to facilitate H3K27me3 deposition, or (ii) ALs counteract PRC2 activity in a chromatin tug-of-war. If ALs acted as PRC2 recruiters, we would expect loss of H3K27me3. However, we did not detect significant loss of genome-wide H3K27me3 occupancy in *al* multiple mutants (Fig. 5c-d). This indicates that AL proteins neither recruit PRC2 nor facilitate its activity.

A second scenario is that ALs and PRC2 engage in a competitive interplay over chromatin states, particularly during early seed development. Similarly to mammals, in flowering plants a large reset of epigenetic information occurs in the gametes and a further lost in the newly formed zygote (Du et al., 2022; Feng et al., 2010; Gutierrez-Marcos and Dickinson, 2012; Jo and Nodine, 2024; Ono and Kinoshita, 2021; Wasserzug-Pash and Klutstein, 2019). This erasure allows the zygote to reinitiate cell division and transition from the highly specialized state of gametes to a totipotent state, enabling the establishment of new cell identities. During this developmental window, chromatin states are being dynamically redefined, making embryonic chromatin highly plastic. In mammals, genome-wide mapping of H3K4me3 and H3K27me3 during early embryogenesis has revealed that borders of these marks are not initially clearly defined, with their resolution increasing progressively through differentiation (Gorkin et al., 2020; Ke et al., 2017; Lu et al., 2016). In *Arabidopsis*, however, exstensive mapping for histone modifications in gametes and early embryos is still lacking.

Nevertheless, given the broad similarities in plant and animal development, we speculate that plants also undergo a comparable epigenetic transition. We hypothesize that during early embryogenesis, PRC2 and AL-containing complexes co-occupy the same or very close genomic regions, competing to define chromatin states. To test this hypothesis, we examined the correlation between H3K4me3 and H3K27me3 in early seeds and compared them to seedlings, a more differentiated tissue (Supplementary Table 3). While in seedlings these two marks are typically well-separated and generally mutually exclusive, early developing seeds exhibited broad overlap between H3K4me3 and H3K27me3 domains with higher correlation (Figs. 5e and EV3b), and thus resembling the less-defined chromatin state observed in early mammalian embryos. This model is further supported by the observation that H3K4me3 loss in *al* mutants is accompanied by a redistribution of H3K27me3, indicating a shift in chromatin boundaries (Fig. 5f and Supplementary Table 3). In the absence of ALs, some genes acquired increased levels of this repressive mark (Fig. 5f and Supplementary Table 3), reinforcing the idea of a dynamic chromatin competition. Integrating histone modification profiles with RNA-Seq data revealed a strong correlation between chromatin state and gene expression: genes that lost H3K4me3 or gained H3K27me3 were downregulated, whereas those that gained H3K4me3 and lost H3K27me3 were upregulated (Fig. EV3c and Supplementary Table 3).

### AL proteins counteract H3K27me3 spreading at developmental genes

Our findings suggest that AL proteins and PRC2 engage in an epigenetic ‘tug-of-war,’ competing to establish chromatin states during early development. In *al* multiple mutants, we identified genes that gain H3K27me3, raising the question of whether this alteration contributes to the developmental defects observed in these plants. To investigate this, we focused on the four *MAF* genes, a clade of *MADS-box* transcription factors that function as repressors of flowering (Ratcliffe et al., 2003). All four *MAF* genes were downregulated in *al3/4/5/6/7* quintuple and *al1/3/4/5/6/7* sextuple mutants (Supplementary Table 2), coinciding with a slight reduction in H3K4me3 levels and a significant increase in H3K27me3, particularly in the sextuple mutant (Fig. 6a and Supplementary Table 3). Notably, these epigenetic changes were concentrated within and around the first long intron of *MAF* genes (Fig. 6a), a known regulatory hotspot for flowering time control (Costa and Dean, 2019; Fu et al., 2005). Flowering repressor genes are typically silenced after a plant has flowered but must be reactivated during embryogenesis to ensure proper flowering regulation in the next generation (Sheldon et al., 2008; Tao et al., 2019). These findings highlight an essential role for AL proteins in balancing histone methylation dynamics to regulate key developmental transitions. To determine whether *MAF* transcription is progressively silenced during embryogenesis, we examined its expression in wild-type and *al1/3/4/5/6/7* mutant embryos. Given that AL proteins do not interfere with de novo H3K4me3 deposition (Fig. 5a-b), we expected *MAF* genes to be initially activated similarly in wild-type and mutant embryos. However, if ALs are required to maintain H3K4me3 occupancy and prevent H3K27me3 accumulation at *MAF* loci, their expression should progressively decline during embryogenesis in *al1/3/4/5/6/7* mutant background, leading to their silencing. Public transcriptome datasets show that *MAF2, MAF3,* and *MAF4* transcripts are detected as early as the 2- to 4-cell embryo stage and follow a reactivation pattern along embryogenesis similar to other flowering repressors, such as *FLOWERING LOCUS C* (*FLC*, (Michaels and Amasino, 1999), Fig. 6b). To test whether *MAF* expression is altered in *al1/3/4/5/6/7* mutants, we performed *in situ* hybridization using a promiscuous antisense probe detecting *MAF1, MAF2,* and *MAF3*. In wild-type embryos, *MAF* expression was uniformly strong at the globular stage, and this pattern remained unchanged in later stages (Fig. 6c). In *al1/3/4/5/6/7* sextuple mutants, *MAFs* expression remained uniform in the early stages even in those embryos that looked slightly malformed (Fig. 6d); however, from the late globular stage onward, mutant embryos exhibited a patchy expression pattern, with some cells expressing *MAFs* at higher or lower levels than others. We interpret this as evidence that, in *al* mutants, individual embryonic cells acquire stochastic epigenetic states, leading to variable *MAFs* expression. We conclude that ALs are required to maintain uniform epigenetic states throughout embryogenesis, preventing the progressive gain of H3K27me3 that ultimately leads to *MAF* silencing. Reduced expression of flowering repressors, including *MAF* genes, is often associated with early flowering phenotypes. To link *MAFs* expression and its embryonic epigenetic pattern to flowering time regulation in adult plants, we examined the flowering time of wild-type and *al1/3/4/5/6/7* mutant plants, also considering that one of the most noticeable macroscopic phenotypes of *al* multiple mutants is indeed an altered flowering time (Fig. E1V).

**Figure 6.**
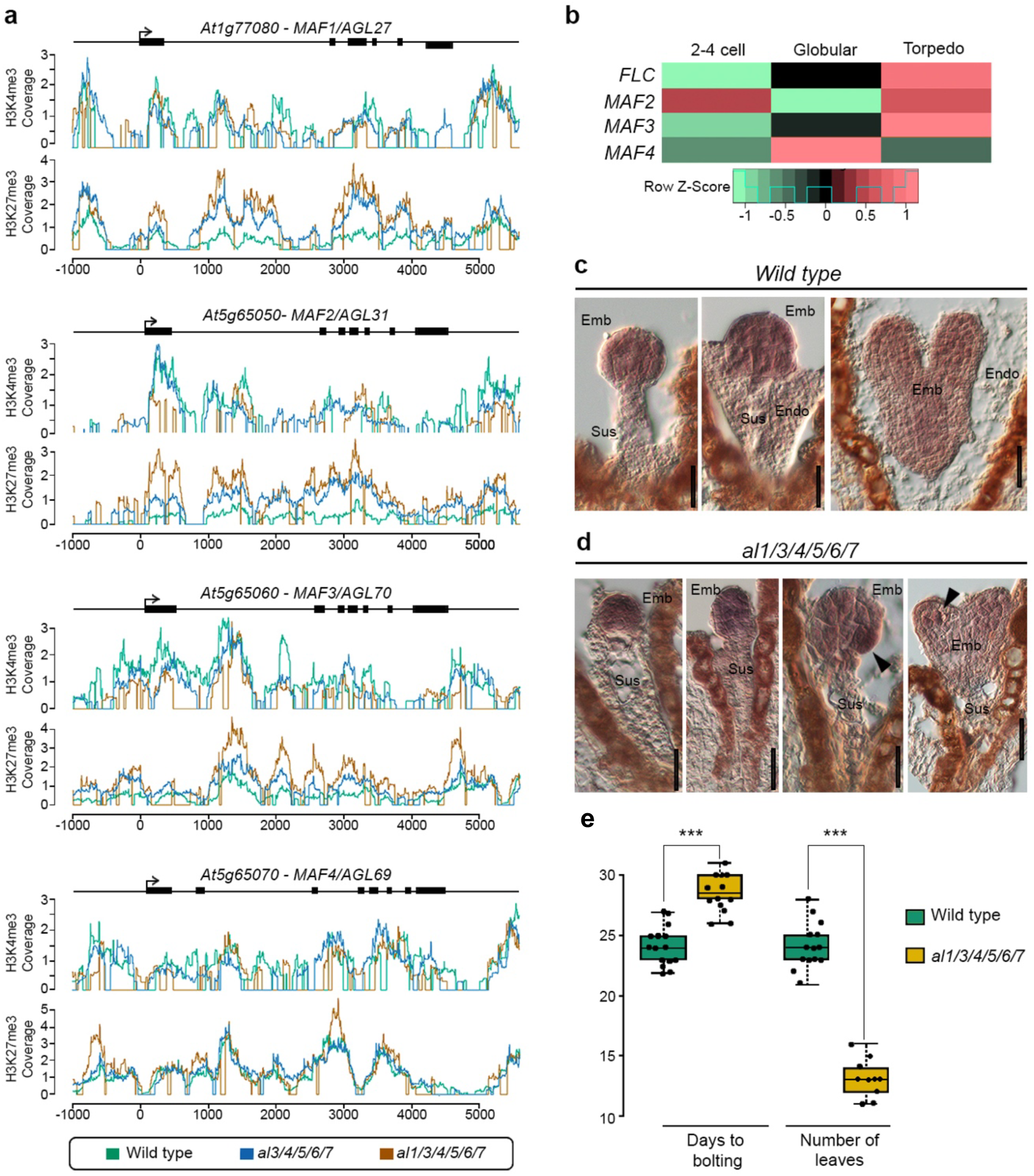
ALs regulate flowering time genes during embryogenesis. **(a)** Histogram showing H3K4me3 and H3K27me3 occupancy at *MAF1, MAF2, MAF3,* and *MAF4* loci in developing seeds of wild-type (green line), *al3/4/5/6/7* quintuple (blue line), and *al1/3/4/5/6/7* sextuple (orange line) mutants. **(b)** Expression levels of *MAF2, MAF3,* and *MAF4* compared to *FLC* during embryogenesis, based on public transcriptome datasets. **(c-d)** *In situ* hybridization using a *MAF* antisense probe in wild-type (c) and *al1/3/4/5/6/7* sextuple (d) embryos from the early globular to heart–torpedo stages. Arrowheads indicate regions in the embryo proper with heterogeneous *MAF* expression. **(e)** Flowering time analysis of wild-type and *al1/3/4/5/6/7* sextuple mutant plants, measured as days to bolting and the number of rosette leaves produced before floral transition. Abbreviations: Emb, embryo; Endo, endosperm; Sus, suspensor. Scale bar: 20 µm.

Flowering time can be assessed by measuring both the number of days required for bolting and the number of leaves produced before the shoot apical meristem transitions from the vegetative to the reproductive phase. In *Arabidopsis*, this transition is unidirectiona (Boss et al., 2004; Sablowski, 2007)l: early flowering mutants initiate reproductive development sooner, producing fewer leaves, whereas late flowering mutants delay the transition and accumulate more leaves before flowering. Since flowering represents an irreversible developmental switch in *Arabidopsis*, a reduced leaf number confirms an early transition to reproductive development, regardless of the absolute time taken to reach that stage. Under our growth conditions, *al1/3/4/5/6/7* sextuple mutants bolted a few days later than wild-type plants, as previously suggested (Su et al., 2025; Xu et al., 2024). However, *al* sextuple mutants produced nearly half as many rosette leaves before bolting compared to wild-type plants (Figure 6f).

This indicates that, despite the apparent bolting delay, the transition to reproductive development occurred earlier. Thus, the observed bolting delay is not due to a prolonged vegetative phase but rather a general reduction in overall growth. We conclude that *al1/3/4/5/6/7* plants exhibit an early flowering phenotype, aligning with our transcriptome and epigenomic analysis.

Altogether, these findings demonstrate that epigenetic states established during embryogenesis persist into adulthood, directly influencing key developmental traits, and position histone modification readers at the center of epigenetic landscape maintenance throughout plant development.

## DISCUSSION

The regulation of gene expression by histone modifications has been extensively studied across different organisms, yet characterizing histone modification readers remains challenging, particularly in plants. Here, we investigated the developmental role of the ALFIN-LIKE (AL) protein family, a class of plant-specific H3K4me3 readers.

AL proteins exhibit broad tissue expression, and previous studies have hinted at their involvement in development. By generating multiple *al* mutants, we uncovered an essential role for ALs during embryogenesis, where they are required to establish cell fate. We detected a range of patterning and growth defects during early embryogenesis, and ALs regulate the expression of several transcription factors with known roles in embryonic development, morphogenesis, and tissue patterning. Interestingly, developmental defects arise predominantly during early embryogenesis. When we analyzed nearly fully developed mutant embryos, the frequency of these defects was significantly reduced. This suggests that the critical errors occur at the earliest developmental stages: once an embryo takes an incorrect trajectory early on, the consequences increase over time, leading to more severe defects. By contrast, embryos that make the correct early developmental decisions proceed normally. This suggests that the interpretation of H3K4me3 is essential for lineage specification and the suppression of suspensor-derived embryogenesis. Similarly, proper embryo patterning depends on AL-mediated chromatin regulation.

One of the most striking phenotypes of *al* mutants is the formation of an extra-embryonic structure growing from the suspensor. This phenotype was first described over two decades ago in pioneering mutagenesis screens aimed at identifying embryo-lethal mutants in *Arabidopsis* (Meinke, 2020). These early studies demonstrated that the suspensor retains embryogenic potential, but that this potential is normally repressed by the embryo proper (Liu et al., 2015). Auxin plays a crucial role in regulating this process (Rademacher et al., 2012), and, notably, gene ontology (GO) terms related to auxin transport and signaling were among the most represented in the transcriptome of *al* multiple mutant seeds.

In both animals and plants, epigenetic information inherited from gametes influences early development. In animals, for instance, H3K27me3 is deposited at specific loci in the oocyte, shaping the initial zygotic transcriptome (Zenk et al., 2017). However, chromatin in the zygote remains highly plastic—a key feature for unlocking totipotency. During early embryogenesis, H3K4me3 is first deposited in the mammalian zygote, followed by H3K27me3, suggesting that the timing and placement of H3K4me3 strongly influence subsequent H3K27me3 deposition (Akkers et al., 2009; Liu et al., 2019). Whether a similar sequence occurs in plants remains unknown. The individual epigenetic landscapes of the *Arabidopsis* egg cell and zygote have yet to be mapped, and the extent to which plant gametes transmit chromatin states to the embryo remains largely unexplored. The technical challenges of isolating plant gametes and profiling chromatin states have so far hindered a detailed understanding of this process. However, analyzing the phenotypes of mutants affecting H3K4me3 and H3K27me3 deposition provides insight into the hierarchy of these modifications. In *Polycomb* group mutants such as *fie* and *mea*, where H3K27me3 deposition is impaired, embryos can still develop until the heart stage, suggesting that early embryogenesis can proceed without H3K27me3 (Bouyer et al., 2011; Grossniklaus et al., 1998; Ohad et al., 1996). By contrast, mutants affecting H3K4me3 deposition, such as *TRAUCO (TRO,* (Aquea et al., 2010)), arrest at the early globular stage, indicating a much earlier role for H3K4me3 in instructing embryogenesis. Our study identifies a similar pattern: the earliest defects in *al* mutants appear immediately after the first zygotic division. Since ALs are H3K4me3 readers, this suggests that H3K4me3 is already present in the zygote epigenome. Whether this mark is inherited from gametes or established *de novo* remains an open question.

Our study reveals an unexpected crosstalk between ALs and PRC2-mediated H3K27me3 deposition which seems particularly relevant for seed development. While chromatin readers have been already implicated in PRC2 recruitment (Godwin and Farrona, 2022), we do not believe this applies to the relationship between ALs and PRC2. Differently, we hypothesize that during early *Arabidopsis* embryogensis, chromatin is highly dynamic, and in this environment, a competing set of histone modification writers and readers engage in a "tug-of-war," to shape the chromatin landscape. The ultimate outcome - whether a gene remains poised for activation or becomes permanently silenced - is determined by the balance of these opposing factors, influenced by developmental cues and chromatin context. These findings highlight the role of AL proteins in reinforcing H3K4me3 deposition, ensuring the stable inheritance of this active chromatin mark and concurrently limiting the spread of repressive H3K27me3. This activity ensures proper gene activation during critical developmental transitions. We present an intriguing example of how epigenetic regulation during embryogenesis can influence traits in the adult plant. The notion that flowering time is preconditioned during embryogenesis is intriguing and has been demonstrated for instance for FLC (Sheldon et al., 2008; Tao et al., 2019). During embryogenesis, *FLC* is reactivated to ensure the plant retains the ability to respond to vernalization later in life. A similar mechanism appears to govern members of the *MAF* gene clade. The stochastic expression of *MAF* genes in *al* multiple mutants resembles the cell-autonomous silencing of *FLC* observed in the *Arabidopsis* root, where each cell independently initiates epigenetic silencing (Berry et al., 2015). Together, these findings underscore the heritable nature of histone modifications and emphasize the critical importance of their correct deposition during the earliest stages of development. Emerging technologies will enable us to dissect how this early epigenetic programming is established during embryogenesis and maintained throughout the plant life cycle, offering new insights into how developmental trajectories are set and preserved over time.

## MATERIAL AND METHODS

### Plant material and growth conditions

Plant material and growth conditions Seeds were sown on half-strength MS media (1/2 MS salt base [Carolina Biologicals, USA], 1% sucrose, 0.05% MES, 0.8% Phytoagar [Duchefa], pH>5.7 with KOH), stratified for 3-4 days at 4°C in the dark, then moved to long-day conditions (8h dark at 18°C, 16h light at 22°C, 70% humidity). At the four-leaf stage, seedlings were transplanted to soil and grown under the same conditions. Marker lines and mutants used: *pCLF::CLF-GFP* in *clf-29*, *pSWN::SWN-GFP* in *swn-4* ((Shu et al., 2019)), *pMEA::MEA-GFP* in *mea-1* (Simonini et al., 2021). AL7 mutant allele (SALK_032503C) was obtained from NASC (described in Jing et al., 2025)). Reciprocal crosses were performed by emasculating stage 11 flowers, allowing 24h for ovule maturation, and manually pollination with the desired pollen.

### CRISPR-Cas9 strategy to create multiple *al* mutants

CRISPR-Cas9 strategy for multiple al mutants Guide RNA sequences for ALFIN-LIKE knockouts were designed using CHOPCHOP v3, selecting three guides per gene. Primers were ordered from IDT and used in PCR with pAGM9037 as the template. Two constructs to mutate AL4 as been assembled following the cloning strategy in (Wang et al., 2015) with single gRNA (AL4-CR862 and AL4-CR902) and used to transform the homozygous mutant for the AL7 gene T1 plants have been selected on Hygromicin. To create a higher level of mutants the gRNA chosen are: 44,45 targeting *AL1*; 46,48 targeting *AL2*; 49,51 targeting *AL3*; 55,57 targeting *AL5*; 59, 60 targeting *AL6*. The gRNAs selected have been assembled following the cloning strategy as in Grüzner et al., 2022 in the final plasmid pAGM65879. T1 plants have been selected for BASTA resistance. A second construct as been created targeting only AL2 using the gRNA 47 and 48 following the strategy for two gRNA plus the selection gene redseed as in Grüzner et al., 2020. T1 plants have been screen for red fluorescence in the seed coat. Each construct was then introduced in Arabidopsis dby Agrobacterium GV3101-mediated transformation.

### Vectors

For construction of the ALs GFP-tagged lines (*pAL3::AL3-GFP*, *pAL4::AL4-GFP* and *pAL7:.AL7-GFP*), their respective genomic sequence were amplified from Col-0 genomic DNA using the couple of primers listed in the Supplementary Table 4.

For *pAL3::AL3-GFP*, the genomic region of the gene *AL3* has been amplified with the primers: olAL3-1 (−4413 from ATG) and AL3-2; olAL3-3b and olAL3-4b to amplify the gene region; olAL3-5b and olAL3-6b to amplify the region 875bp downstream the gene. Fragments were subcloned in pJET1.2 (Thermoscientific) and assembled together with the GFP plasmid pSB279 in the backbone pSB280 with the enzyme SapI following Golden Gate cloning strategy.

For *pAL4::AL4-GFP*, the genomic region of the gene AL4 has been amplified with the primers: olAL4-1 (−2979 from ATG) and AL4-2; olAL4-3 and olAL4-4 to amplify the gene region; olAL4-5 and olAL4-6 to amplify the region 294 downstream the gene. Fragments were subcloned in pJET1.2 (Thermoscientific) and assembled together with the GFP plasmid pICSL50008 in the backbone pSTB140 with the enzyme BsaI following Golden Gate cloning strategy.

For pAL7::AL7-GFP,

The PCR fragments were cloned into the miniT vector (NEB), and assembled together in a Golden Gate reaction together with the terminator tHSP18.2 (Addgene GB0035) and the destination vector p140 using the enzyme BsaI. The destination vector p140 is a modified version of the Golden Gate low copy binary vector pAGM53451(Grützner et al., 2021). For our purpose, we inserted in the pAGM53451 vector the RedSeed selector marker at position three, whereas at position two we inserted the lacZ gene adapted to be an acceptor for Level 1 Golden Gate modules. The final constructs has been transformed in *A.tumefaciens* GV3101 by floral-dip (Clough and Bent, 1998) and then in the *al1/3/4/5/6/7* sextuple mutant. Plants that showed recovered phenotypes like the *al1/3/5/6/7* quintuple mutant were selected and brought to homozygosity.

### Clearing

Siliques were fixed overnight in ethanol:acetic acid (9:1), transferred to 70% ethanol, and dissected. Seeds/ovules were cleared in chloral hydrate solution for a few hours or overnight and imaged using a Zeiss DMR microscope with DIC filters and a Leica Flexacam C3 LSR camera.

### Imaging of GFP-tagged lines

ALs-GFP marker lines were imaged with a Leica TCS SP8 Multiphoton microscope in photon counting mode. Ovules and seeds were mounted in 7% glucose with 10 µg/mL Propidium Iodide.

### *In situ* hybridization

The digoxygenin-labelled antisense RNA probes for WOX2 and MAFs were generated by *in vitro* transcription according to the instructions provided with the DIG RNA labelling kit (SP6/T7; Roche) using a plasmid containing the individual probes sequence as template (For sequences, see Supplementary Table 4). Developing siliques were collected at the desired time points and fixed in FAA fixative (Formaldehyde, acetic acid, ethanol) o/n, then moved in ethanol 70% and into an ASP200 embedding machine for embedding (Leica Microsystems GmbH, Wetzlar, Germany). The samples were then sliced in 8-μm sections with a RM2145 Leica microtome (Leica Microsystems GmbH, Wetzlar, Germany) and then hybridized as described previously, with strong formaldehyde washes, as described in (Dreni et al., 2007). Images were taken with a Zeiss DMR microscope equipped with differential interference contrast (DIC) filters and Leica Flexacam C3 LSR camera.

### Protein immunoprecipitation coupled with liquid chromatography tandem mass spectrometry (LC-MS/MS)

For immunoprecipitation followed by liquid chromatography-tandem mass spectrometry (IP LC-MS/MS) experiments, *pCLF::CLF-GFP* and *pSWN::SWN-GFP* seedlings at 14 days post-germination were harvested and stored at −80°C until further processing. For IP LC-MS/MS experiments involving *pCLF::CLF-GFP*, *pSWN::SWN-GFP*, *pMEA::MEA-GFP*, and *pAL4::AL4-GFP* lines, inflorescence tissue and flowers at 1-2 days after anthesis were collected, whereas siliques at 1-4 days after fertilization were collected from *pAL4::AL4-GFP* plants. All samples were flash-frozen in liquid nitrogen and stored at −80°C until further use. For the immunoprecipitation step, we followed the protocol reported in Wendrich et al., 2017. The frozen tissue was ground in liquid nitrogen for 30 minutes, and the resulting powder was collected in a 50 mL Falcon tube and stored at −80°C. The following day, the frozen powder was resuspended in EB+ buffer (50 mM Tris-HCl, pH 7.5, 150 mM NaCl, 1% NP-40, protease inhibitor pellet) at a ratio of approximately 30 mL EB+ buffer per 10 g of powder. The homogenate was filtered through Miracloth (Merck KGaA, Darmstadt) and sonicated using a Bioruptor (five cycles of 15 s on and 15 s off, medium power). Samples were incubated on ice for 30 minutes before centrifugation at 39,000g for 15 minutes at 4°C to remove debris. The supernatant was further filtered using a 40 µm cell strainer. For immunoprecipitation, anti-GFP beads were added at a ratio of 1:500 and incubated with rotation at 4°C for 2 hours. GFP-Trap Magnetic Agarose (Chromotek) were used for CLF-GFP, SWN-GFP, and AL4-GFP immunoprecipitations. Anti-GFP MicroBeads (Miltenyi Biotec) were used for MEA-GFP samples. Immunoprecipitations were performed according to the manufacturer’s protocol. Beads were washed with EB+ buffer and eluted in ABC buffer (50 mM NH_4_HCO_3_). Eluted samples were further processed and prepared for LC-MS/MS analysis at the Functional Genomics Center Zurich starting with an On-beads digestion with Trypsin, and precipitation of the peptides with 0.1% trifluoroacetic acid (TFA) in 50% acetonitrile. All fractions were combined and dried before further processing. Peptide concentrations were estimated using the Lunatic UV/Vis absorbance spectrometer (Unchained Lab). Digested peptide samples were dissolved in 3% acetonitrile with 0.1% formic acid and analyzed using a liquid chromatography-tandem mass spectrometry (LC-MS/MS) workflow. Peptides were separated on an M-class Ultra-Performance Liquid Chromatography (UPLC) system and analyzed using an Orbitrap mass spectrometer (Thermo Scientific). Peptide concentrations were re-evaluated using the Lunatic UV/Vis absorbance spectrometer prior to LC-MS injection. Basic quality control of raw LC-MS data, including total ion chromatogram (TIC), base peak chromatogram (BPC), lock-mass correction, and internal retention time (iRT) peptide signal assessment, was performed using the rawDiag-shiny application (Trachsel et al., 2018).

Label-free quantification (LFQ) with match-between-runs (MBR) was performed using the FragPipe-RESOURCE workflow executed within the b-fabric app 295. Data processing was conducted using the Philosopher command-line interface (CLI, (da Veiga Leprevost et al., 2020)), and LFQ was carried out with the IonQuant software tool (Yu et al., 2021).

### RNA-Seq

The RNA-Seq has been performed on isolated seeds from 1 to 4 days after fertilization. Siliques were fixed on a double-sided tape, open along the replum with a fine needle, seeds gently scraped off with a needle and immediately placed in an Eppendorf filled with RNAlater buffer (ThermoFisher) and placed on ice. After maximum one hour of collection, seeds were precipitated by centrifugating the tubes at 500g for 2min at RT, the supernatant removed, and the seed pellet stored at −70 until the day of extraction. Total RNA was extracted using the Vazyme FastPure Universal Plant Total RNA Isolation Kit, following the manufacturer’s instructions. RNA integrity and concentration were assessed using the TapeStation 4200 and RNA ScreenTape Analysis. DNA contamination was removed by treatment with DNase I (Invitrogen™ TURBO DNase™). First-strand cDNA synthesis was performed using Invitrogen™ SuperScript™ II Reverse Transcriptase with oligo(dT) primers and 5 µg of total RNA. Second-strand synthesis followed the Gubler & Hoffman method (Gubler and Hoffman, 1983). Strand-specific libraries were prepared using the Nextera XT DNA Kit. Sequencing was carried out at the Functional Genomics Center Zurich (FGCZ) on an Illumina NovaSeq X Plus 1 Lane 10B Flowcell. cDNA and concentration were assessed using the TapeStation 4200 and DNA and High Sensitivity ScreenTape Analysis. All bioinformatics analyses were conducted on a local machine running Ubuntu 24.04.1 LTS (Noble Numbat) within a Conda 24.9.2 environment. Quality control was performed with FastQC v0.12.1 and MultiQC v1.25.2, while adapter trimming was conducted using Fastp v0.24.0 (Chen et al., 2018). Read mapping and gene expression quantification were performed with STAR v2.7.11b (Dobin et al., 2013). The *Arabidopsis thaliana* reference genome used was Araport11 (Cheng et al., 2017). Differential expression analysis was conducted in R v4.4.3 using DESeq2 v1.46.0, applying a log2FC > 0.5 and an adjusted p-value < 0.01 as significance thresholds. Gene ontology (GO) term enrichment analysis was performed using PLAZA 5.0 (Van Bel et al., 2022). DE genes were clustered by hierarchical clustering using z-scores with an Euclidean distance and Wards’s agglomeration method.

### CUT&Tag

For the CUT&Tag, we followed the protocol described in Fu et al., 2024 Biorxiv (Fu et al., 2024). Antibodies used in this study are: Rabbit anti-H3K27me3 (Cell signaling technology #9733, dilution ratio 1:50); Rabbit anti H3K4me3 (Diagenode c15410003, dilution ratio 1:150); Mouse anti-H3 (Active Motif 39064, dilution ratio 1:150). The generated libraries were pooled and sequenced on an Illumina NovaSeq X Plus sequencer. For bioinformatic data processing, short reads generated in this study were deposited at NCBI Sequence Read Archive (SRA, www.ncbi.nlm.nih.gov/sra) and are accessible through the accession number XXX. Following removal of sequencing adapters and low-quality bases with fastp (version 0.23.4 with the options −l 30 -a CTGTCTCTTATACACATCT) paired-end reads were aligned to the Arabidopsis thaliana reference genome (TAIR10) using bowtie2 (version 2.4.5 with the options --local --very-sensitive --no-mixed --no-unal --no-discordant --phred33 -I 10 -X 700, (Langmead and Salzberg, 2012)). Only reads with a minimal alignment quality of 10 were kept. Duplicate reads were removed with Picard tools (version 3.1.1, broadinstitute.github.io/picard). Peak calling was done with MACS2 using the matching H3 libraries as background control (version 2.2.7.1 with the options -f BAMPE -g 119146348 -- nomodel -q 0.05 --broad --broad-cutoff 0.1, (Zhang et al., 2008)). Peaks different replicates were merged with multovl (version 1.3 with the option -m 1 -u, (Aszódi, 2012)) and reads per peak were counted with featureCounts (version 2.0.6 with the options -O -p, (Liao et al., 2014)). Peaks were intersected with the Arabidopsis thaliana gene annotation (Araport11) using 3/1.5 kb up-/down-stream flanking sequences and the program mapGenomicRangesOverlap (github.com/MWSchmid/mapGenomicRanges).

For differential binding analysis, variation in counts per peak was analyzed with a general linear model in R with the package DESeq (version 1.38.3; (Love et al., 2014)) according to a factorial design with all two explanatory factors combined into a single factor (Schmid, 2017). Specific conditions were compared with linear contrasts. Within each comparison, p-values were adjusted for multiple testing (Benjamini-Hochberg), and peaks with an adjusted p-value (false discovery rate, FDR) below 0.05 and a minimal log2 fold-change (i.e., the difference between the log2 transformed, normalized sequence counts) of 1.5 were considered to be differentially bound. Normalized sequence counts were calculated accordingly with DESeq2 and log2(x+1) transformed.

For functional characterization (Gene Ontology), we tested for enrichment of gene ontology (GO) terms using topGO (version 2.50, (Alexa et al., 2006)) in conjunction with the GO annotation available through biomaRt (Durinck et al., 2009). Analysis was based on gene counts using the "weight" algorithm with Fisher’s exact test (both implemented in topGO). A term was identified as significant if the p-value was below 0.01. For data visualizations (to visualize coverage along a genomic stretch), coverage was depth-normalized with bedtools genomecov (version 2.31.0, (Quinlan and Hall, 2010)) and BedGraphs were converted to BigWigs with bedGraphToBigWig (version 2.10, (Kent et al., 2010)). Coverage heatmaps were drawn using deeptools as described in the tutorial (version 3.5.1, (Ramírez et al., 2014)). If more than 1,000 were available for display, sites were downsampled randomly to 1,000 sites.

## SUPPLEMENTAL INFORMATION

Supplemental Information can be found online at …

## ACKNOWLEDGEMENTS

We thank Yuhai Cui (Agriculture and Agri-Food Canada), and the Nottingham Arabidopsis Stock Center for providing seeds, Stefano Bencivenga and Max Schwarze (University of Zurich) for helpful discussion or comments on the manuscript, Diana Zörner, Christof Eichenberger, Arturo Bolaños, and Agnesa Lleshaj-Pitaqi (University of Zurich) for general lab support, and the Functional Genomic Centre Zurich for technical assistance. We thank Ueli Grossniklaus (University of Zurich), in whose laboratory this project was initiated, and for generously sharing unpublished data. This work was supported by the University of Zurich, and grants from the Swiss National Science Foundation to SS.

## AUTHOR CONTRIBUTIONS

JS, YF, ET performed experiments, MWS did the bioinformatic analysis, SS conceived, supervised, raised funding for the project, performed experiments and wrote the manuscript with input from all the authors.

## DATA AVAILABILITY

The raw RNA-Seq and CUT&Tag dataset generated in this work can be accessed at XXX.

## Supplementary Material

**Figure EV1.**
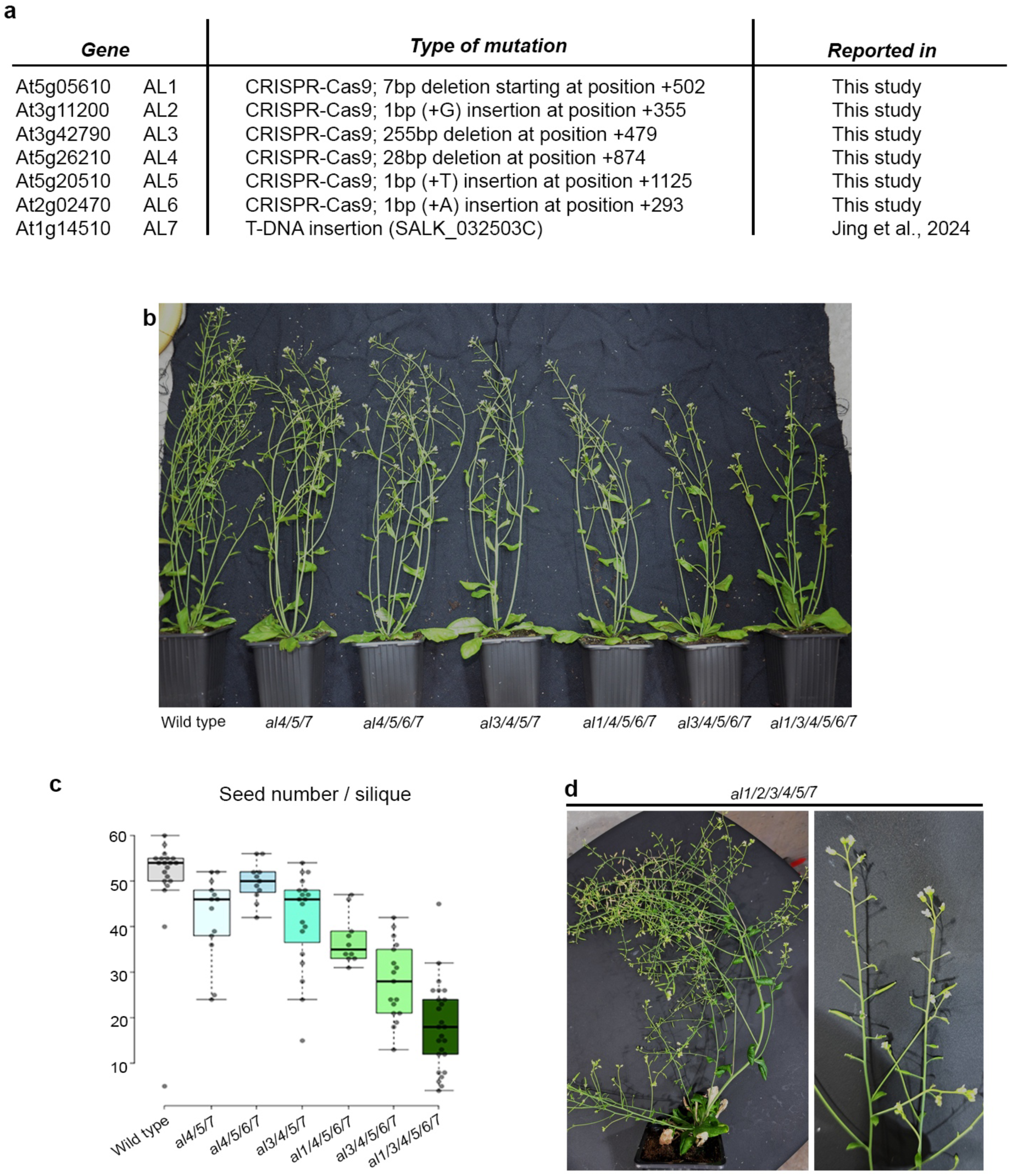
Macroscopic morphological analysis of *al* multiple mutants generated using CRISPR-Cas9. **(a)** Summary of CRISPR-Cas9-induced alleles obtained for *AL1, AL2, AL3, AL4, AL5,* and *AL6*. **(b)** Overall phenotype of *al* multiple mutant plants. **(c)** Seed set of *al* multiple mutant plants, represented as the number of seeds per silique. **(d)** Overall phenotype of *al2/3/4/5/6/7* sextuple mutant plants.

**Figure EV2.**
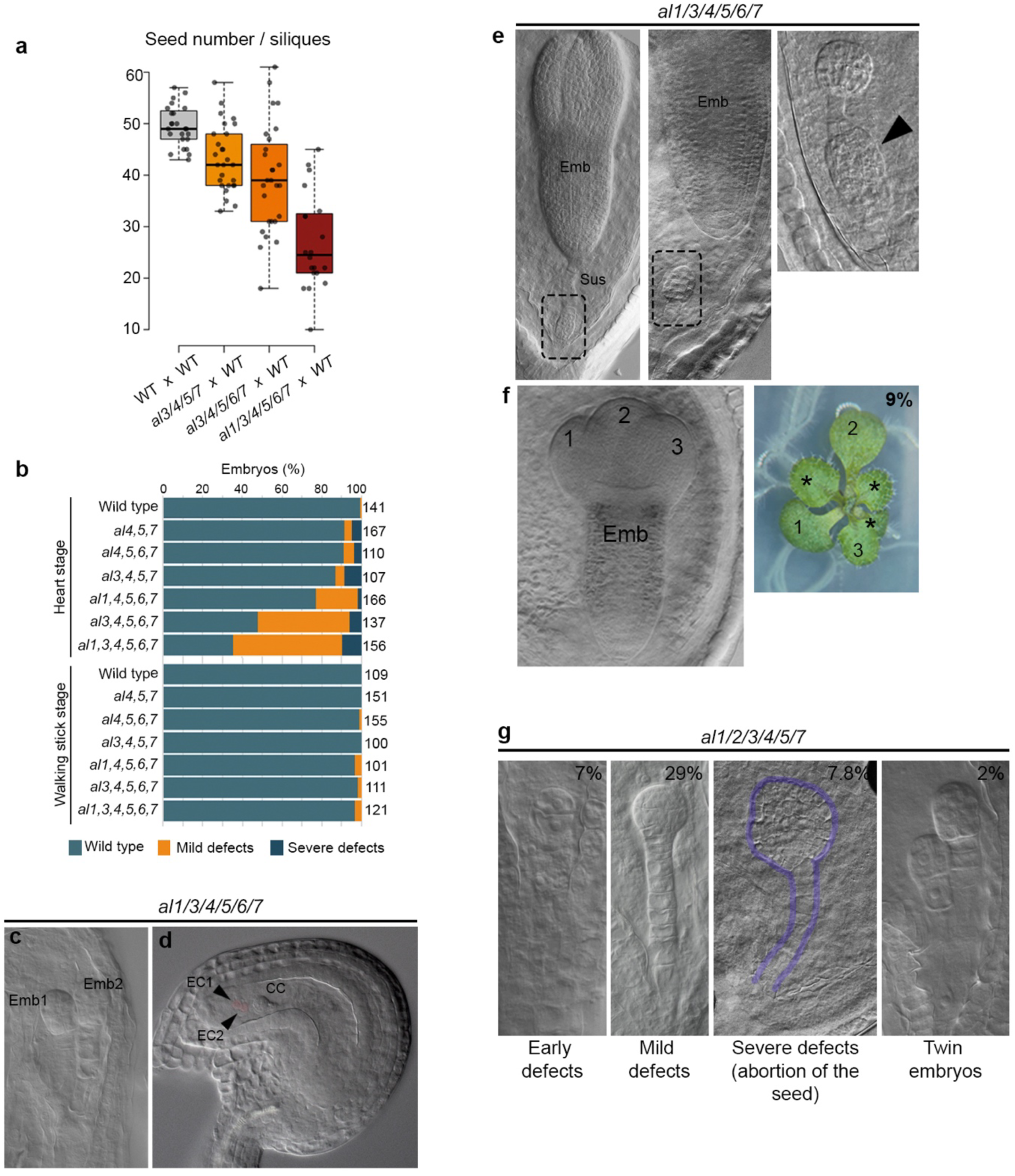
Developmental defects of *al* multiple mutants during the reproductive phase and seed development. **(a)** Reciprocal crosses between *al* multiple mutants and wild-type pollen to assess potential female gametophytic defects, represented as the number of seeds per silique. **(b)** Quantification of embryonic phenotypes (wild-type-like, mild, and severe developmental defects) in different *al* mutant combinations at the heart and walking-stick embryonic stages. **(c)** Twin embryos developing within a single embryo sac in the *al1/3/4/5/6/7* sextuple mutant, with minimal surrounding endosperm. **(d)** Clearing image of an *al1/3/4/5/6/7* ovule exhibiting two egg cells, artificially colored in magenta. **(e)** Extra-embryonic embryo developing from the suspensor in *al1/3/4/5/6/7* sextuple mutants. **(f)** Embryos and seedlings with three cotyledons in the *al1/3/4/5/6/7* sextuple mutant. The percentage of seedlings exhibiting this phenotype is shown in the top right corner. **(g)** Embryonic defects observed in the *al2/3/4/5/6/7* sextuple mutant, with their quantification (percentage) displayed in the top right corner.

**Figure EV3.**
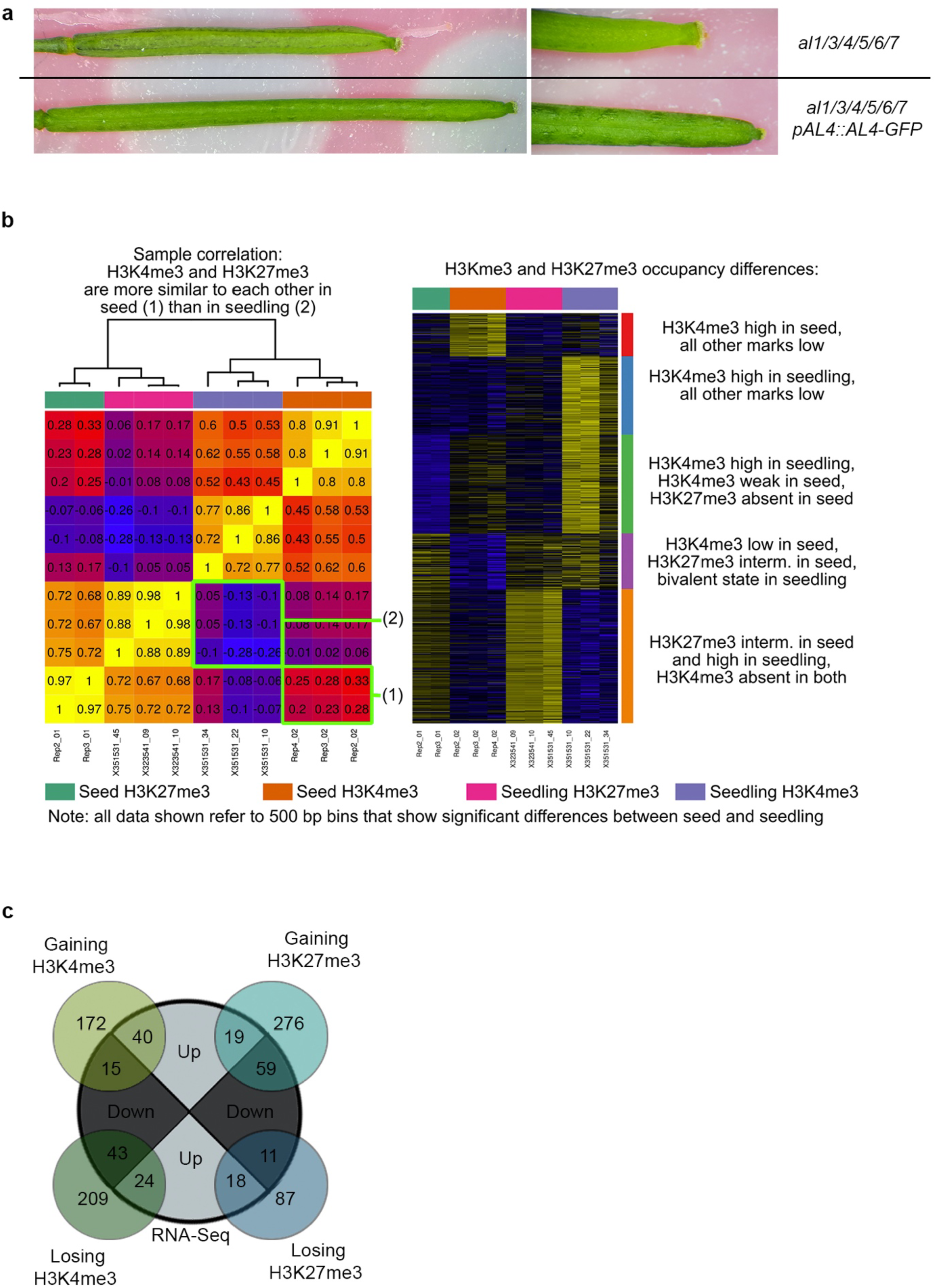
Interaction-proteomics, trasncriptomic and epigenomic analysis. **(a)** *pAL4:.AL4-GFP* transgene complements fruit length and pistil defects of the *al1/3/4/5/6/7* mutant, confirming that the AL4-GFP protein is fully functional. **(b)** Correlation matrix and heat map of H3K27me3 and H3K4me3 occupancy in wild-typeseeds and seedlings. **(c)** Overlap of CUT&Tag and RNA-Seq data, showing correlation between the loss/gain of a given histone mark and the expression level.

## REFERENCES

Akkers, R.C., van Heeringen, S.J., Jacobi, U.G., Janssen-Megens, E.M., Françoijs, K.-J., Stunnenberg, H.G., Veenstra, G.J.C., 2009. A hierarchy of H3K4me3 and H3K27me3 acquisition in spatial gene regulation in Xenopus embryos. Dev. Cell 17, 425–434. 10.1016/j.devcel.2009.08.005

Alexa, A., Rahnenführer, J., Lengauer, T., 2006. Improved scoring of functional groups from gene expression data by decorrelating GO graph structure. Bioinforma. Oxf. Engl. 22, 1600–1607. 10.1093/bioinformatics/btl140

Alvarez-Venegas, R., Pien, S., Sadder, M., Witmer, X., Grossniklaus, U., Avramova, Z., 2003. ATX-1, an Arabidopsis homolog of trithorax, activates flower homeotic genes. Curr. Biol. CB 13, 627– 637. 10.1016/s0960-9822(03)00243-4

Aquea, F., Johnston, A.J., Cañon, P., Grossniklaus, U., Arce-Johnson, P., 2010. TRAUCO, a Trithorax-group gene homologue, is required for early embryogenesis in Arabidopsis thaliana. J. Exp. Bot. 61, 1215–1224. 10.1093/jxb/erp396

Aszódi, A., 2012. MULTOVL: fast multiple overlaps of genomic regions. Bioinforma. Oxf. Engl. 28, 3318–3319. 10.1093/bioinformatics/bts607

Baile, F., Merini, W., Hidalgo, I., Calonje, M., 2020. Dissection of PRC1 and PRC2 recruitment in Arabidopsis connects EAR repressome to PRC2 anchoring (preprint). Plant Biology. 10.1101/2020.08.28.271999

Berger, S.L., 2007. The complex language of chromatin regulation during transcription. Nature 447, 407–412. 10.1038/nature05915

Berry, S., Hartley, M., Olsson, T.S.G., Dean, C., Howard, M., 2015. Local chromatin environment of a Polycomb target gene instructs its own epigenetic inheritance. eLife 4, e07205. 10.7554/eLife.07205

Boss, P.K., Bastow, R.M., Mylne, J.S., Dean, C., 2004. Multiple Pathways in the Decision to Flower: Enabling, Promoting, and Resetting. Plant Cell 16, S18–S31. 10.1105/tpc.015958

Bouyer, D., Roudier, F., Heese, M., Andersen, E.D., Gey, D., Nowack, M.K., Goodrich, J., Renou, J.-P., Grini, P.E., Colot, V., Schnittger, A., 2011. Polycomb Repressive Complex 2 Controls the Embryo-to-Seedling Phase Transition. PLoS Genet. 7, e1002014. 10.1371/journal.pgen.1002014

Chen, S., Zhou, Y., Chen, Y., Gu, J., 2018. fastp: an ultra-fast all-in-one FASTQ preprocessor. Bioinforma. Oxf. Engl. 34, i884–i890. 10.1093/bioinformatics/bty560

Cheng, C.-Y., Krishnakumar, V., Chan, A.P., Thibaud-Nissen, F., Schobel, S., Town, C.D., 2017. Araport11: a complete reannotation of the Arabidopsis thaliana reference genome. Plant J. Cell Mol. Biol. 89, 789–804. 10.1111/tpj.13415

Clough, S.J., Bent, A.F., 1998. Floral dip: a simplified method forAgrobacterium-mediated transformation ofArabidopsis thaliana: Floral dip transformation of Arabidopsis. Plant J. 16, 735–743. 10.1046/j.1365-313x.1998.00343.x

Costa, S., Dean, C., 2019. Storing memories: the distinct phases of Polycomb-mediated silencing of *Arabidopsis FLC*. Biochem. Soc. Trans. 47, 1187–1196. 10.1042/BST20190255

da Veiga Leprevost, F., Haynes, S.E., Avtonomov, D.M., Chang, H.-Y., Shanmugam, A.K., Mellacheruvu, D., Kong, A.T., Nesvizhskii, A.I., 2020. Philosopher: a versatile toolkit for shotgun proteomics data analysis. Nat. Methods 17, 869–870. 10.1038/s41592-020-0912-y

De Lucia, F., Crevillen, P., Jones, A.M.E., Greb, T., Dean, C., 2008. A PHD-polycomb repressive complex 2 triggers the epigenetic silencing of FLC during vernalization. Proc. Natl. Acad. Sci. U. S. A. 105, 16831–16836. 10.1073/pnas.0808687105

Dobin, A., Davis, C.A., Schlesinger, F., Drenkow, J., Zaleski, C., Jha, S., Batut, P., Chaisson, M., Gingeras, T.R., 2013. STAR: ultrafast universal RNA-seq aligner. Bioinforma. Oxf. Engl. 29, 15–21. 10.1093/bioinformatics/bts635

Dreni, L., Jacchia, S., Fornara, F., Fornari, M., Ouwerkerk, P.B.F., An, G., Colombo, L., Kater, M.M., 2007. The D-lineage MADS-box gene OsMADS13 controls ovule identity in rice. Plant J. Cell Mol. Biol. 52, 690–699. 10.1111/j.1365-313X.2007.03272.x

Du, Z., Zhang, K., Xie, W., 2022. Epigenetic Reprogramming in Early Animal Development. Cold Spring Harb. Perspect. Biol. 14, a039677. 10.1101/cshperspect.a039677

Durinck, S., Spellman, P.T., Birney, E., Huber, W., 2009. Mapping identifiers for the integration of genomic datasets with the R/Bioconductor package biomaRt. Nat. Protoc. 4, 1184–1191. 10.1038/nprot.2009.97

Farrona, S., Hurtado, L., March-Díaz, R., Schmitz, R.J., Florencio, F.J., Turck, F., Amasino, R.M., Reyes, J.C., 2011. Brahma is required for proper expression of the floral repressor FLC in Arabidopsis. PloS One 6, e17997. 10.1371/journal.pone.0017997

Feng, S., Jacobsen, S.E., Reik, W., 2010. Epigenetic reprogramming in plant and animal development. Science 330, 622–627. 10.1126/science.1190614

Fu, D., Szűcs, P., Yan, L., Helguera, M., Skinner, J.S., von Zitzewitz, J., Hayes, P.M., Dubcovsky, J., 2005. Large deletions within the first intron in VRN-1 are associated with spring growth habit in barley and wheat. Mol. Genet. Genomics 273, 54–65. 10.1007/s00438-004-1095-4

Fu, Y., Schmid, M.W., Simonini, S., 2024. CUT&amp;Tag for high-resolution epigenomic profiling from a low amount of Arabidopsis tissue. 10.1101/2024.07.29.604300

Gao, Z., Li, Y., Ou, Y., Yin, M., Chen, T., Zeng, X., Li, R., He, Y., 2023. A pair of readers of bivalent chromatin mediate formation of Polycomb-based “memory of cold” in plants. Mol. Cell 83, 1109–1124.e4. 10.1016/j.molcel.2023.02.014

Gendall, A.R., Levy, Y.Y., Wilson, A., Dean, C., 2001. The VERNALIZATION 2 gene mediates the epigenetic regulation of vernalization in Arabidopsis. Cell 107, 525–535. 10.1016/s0092-8674(01)00573-6

Godwin, J., Farrona, S., 2022. The Importance of Networking: Plant Polycomb Repressive Complex 2 and Its Interactors. Epigenomes 6, 8. 10.3390/epigenomes6010008

Gorkin, D.U., Barozzi, I., Zhao, Y., Zhang, Y., Huang, H., Lee, A.Y., Li, B., Chiou, J., Wildberg, A., Ding, B., Zhang, B., Wang, M., Strattan, J.S., Davidson, J.M., Qiu, Y., Afzal, V., Akiyama, J.A., Plajzer-Frick, I., Novak, C.S., Kato, M., Garvin, T.H., Pham, Q.T., Harrington, A.N., Mannion, B.J., Lee, E.A., Fukuda-Yuzawa, Y., He, Y., Preissl, S., Chee, S., Han, J.Y., Williams, B.A., Trout, D., Amrhein, H., Yang, H., Cherry, J.M., Wang, W., Gaulton, K., Ecker, J.R., Shen, Y., Dickel, D.E., Visel, A., Pennacchio, L.A., Ren, B., 2020. An atlas of dynamic chromatin landscapes in mouse fetal development. Nature 583, 744–751. 10.1038/s41586-020-2093-3

Grossniklaus, U., Paro, R., 2014. Transcriptional silencing by polycomb-group proteins. Cold Spring Harb. Perspect. Biol. 6, a019331. 10.1101/cshperspect.a019331

Grossniklaus, U., Vielle-Calzada, J.P., Hoeppner, M.A., Gagliano, W.B., 1998. Maternal control of embryogenesis by MEDEA, a polycomb group gene in Arabidopsis. Science 280, 446–450. 10.1126/science.280.5362.446

Grützner, R., Martin, P., Horn, C., Mortensen, S., Cram, E.J., Lee-Parsons, C.W.T., Stuttmann, J., Marillonnet, S., 2021. High-efficiency genome editing in plants mediated by a Cas9 gene containing multiple introns. Plant Commun. 2, 100135. 10.1016/j.xplc.2020.100135

Gubler, U., Hoffman, B.J., 1983. A simple and very efficient method for generating cDNA libraries. Gene 25, 263–269. 10.1016/0378-1119(83)90230-5

Gutierrez-Marcos, J.F., Dickinson, H.G., 2012. Epigenetic Reprogramming in Plant Reproductive Lineages. Plant Cell Physiol. 53, 817–823. 10.1093/pcp/pcs052

Huanca-Mamani, W., Garcia-Aguilar, M., León-Martínez, G., Grossniklaus, U., Vielle-Calzada, J.-P., 2005. CHR11, a chromatin-remodeling factor essential for nuclear proliferation during female gametogenesis in Arabidopsis thaliana. Proc. Natl. Acad. Sci. U. S. A. 102, 17231–17236. 10.1073/pnas.0508186102

Hyun, K., Jeon, J., Park, K., Kim, J., 2017. Writing, erasing and reading histone lysine methylations. Exp. Mol. Med. 49, e324–e324. 10.1038/emm.2017.11

Ingham, P.W., 1983. Differential expression of bithorax complex genes in the absence of the extra sex combs and trithorax genes. Nature 306, 591–593. 10.1038/306591a0

Jin, R., Yang, H., Muhammad, T., Li, X., Tuerdiyusufu, D., Wang, B., Wang, J., 2024. Involvement of Alfin-Like Transcription Factors in Plant Development and Stress Response. Genes 15, 184. 10.3390/genes15020184

Jing, H., Liu, W., Qu, G.-P., Niu, D., Jin, J.B., 2025. SUMOylation of AL6 regulates seed dormancy and thermoinhibition in Arabidopsis. New Phytol. 245, 1040–1055. 10.1111/nph.20270

Jo, L., Nodine, M.D., 2024. “To remember or forget: Insights into the mechanisms of epigenetic reprogramming and priming in early plant embryos.” Curr. Opin. Plant Biol. 81, 102612. 10.1016/j.pbi.2024.102612

Kayum, M.A., Park, J.-I., Ahmed, N.U., Jung, H.-J., Saha, G., Kang, J.-G., Nou, I.-S., 2015. Characterization and stress-induced expression analysis of Alfin-like transcription factors in Brassica rapa. Mol. Genet. Genomics MGG 290, 1299–1311. 10.1007/s00438-015-0993-y

Ke, Y., Xu, Y., Chen, X., Feng, S., Liu, Z., Sun, Y., Yao, X., Li, F., Zhu, W., Gao, L., Chen, H., Du, Z., Xie, W., Xu, X., Huang, X., Liu, J., 2017. 3D Chromatin Structures of Mature Gametes and Structural Reprogramming during Mammalian Embryogenesis. Cell 170, 367–381.e20. 10.1016/j.cell.2017.06.029

Kent, W.J., Zweig, A.S., Barber, G., Hinrichs, A.S., Karolchik, D., 2010. BigWig and BigBed: enabling browsing of large distributed datasets. Bioinforma. Oxf. Engl. 26, 2204–2207. 10.1093/bioinformatics/btq351

Kim, D.-H., Sung, S., 2017. The Binding Specificity of the PHD-Finger Domain of VIN3 Moderates Vernalization Response1[OPEN]. Plant Physiol. 173, 1258–1268. 10.1104/pp.16.01320

Kiyosue, T., Ohad, N., Yadegari, R., Hannon, M., Dinneny, J., Wells, D., Katz, A., Margossian, L., Harada, J.J., Goldberg, R.B., Fischer, R.L., 1999. Control of fertilization-independent endosperm development by the *MEDEA* polycomb gene in *Arabidopsis*. Proc. Natl. Acad. Sci. 96, 4186–4191. 10.1073/pnas.96.7.4186

Köhler, C., Hennig, L., Bouveret, R., Gheyselinck, J., Grossniklaus, U., Gruissem, W., 2003. Arabidopsis MSI1 is a component of the MEA/FIE Polycomb group complex and required for seed development. EMBO J. 22, 4804–4814. 10.1093/emboj/cdg444

Lai, Y., Lu, X.M., Daron, J., Pan, S., Wang, J., Wang, W., Tsuchiya, T., Holub, E., McDowell, J.M., Slotkin, R.K., Roch, K.G.L., Eulgem, T., 2020. The Arabidopsis PHD-finger protein EDM2 has multiple roles in balancing NLR immune receptor gene expression. PLOS Genet. 16, e1008993. 10.1371/journal.pgen.1008993

Langmead, B., Salzberg, S.L., 2012. Fast gapped-read alignment with Bowtie 2. Nat. Methods 9, 357–359. 10.1038/nmeth.1923

Liang, X., Lei, M., Li, F., Yang, X., Zhou, M., Li, B., Cao, Y., Gong, S., Liu, K., Liu, J., Qi, C., Liu, Y., 2018. Family-Wide Characterization of Histone Binding Abilities of PHD Domains of AL Proteins in Arabidopsis thaliana. Protein J. 37, 531–538. 10.1007/s10930-018-9796-4

Liao, Y., Smyth, G.K., Shi, W., 2014. featureCounts: an efficient general purpose program for assigning sequence reads to genomic features. Bioinforma. Oxf. Engl. 30, 923–930. 10.1093/bioinformatics/btt656

Liu, J., An, L., Wang, J., Liu, Z., Dai, Y., Liu, Y., Yang, L., Du, F., 2019. Dynamic patterns of H3K4me3, H3K27me3, and Nanog during rabbit embryo development. Am. J. Transl. Res. 11, 430–441.

Liu, Y., Li, X., Zhao, J., Tang, X., Tian, S., Chen, J., Shi, C., Wang, W., Zhang, L., Feng, X., Sun, M.-X., 2015. Direct evidence that suspensor cells have embryogenic potential that is suppressed by the embryo proper during normal embryogenesis. Proc. Natl. Acad. Sci. U. S. A. 112, 12432–12437. 10.1073/pnas.1508651112

Long, J.A., Ohno, C., Smith, Z.R., Meyerowitz, E.M., 2006. TOPLESS regulates apical embryonic fate in Arabidopsis. Science 312, 1520–1523. 10.1126/science.1123841

Love, M.I., Huber, W., Anders, S., 2014. Moderated estimation of fold change and dispersion for RNA-seq data with DESeq2. Genome Biol. 15, 550. 10.1186/s13059-014-0550-8

Lu, F., Liu, Y., Inoue, A., Suzuki, T., Zhao, K., Zhang, Y., 2016. Establishing Chromatin Regulatory Landscape during Mouse Preimplantation Development. Cell 165, 1375–1388. 10.1016/j.cell.2016.05.050

Meagher, R.B., Kandasamy, Muthugapatti K., Smith, Aaron P., and McKinney, E.C., 2010. Nuclear actin-related proteins at the core of epigenetic control. Plant Signal. Behav. 5, 518–522. 10.4161/psb.10986

Meinke, D.W., 2020. Genome-wide identification of EMBRYO-DEFECTIVE (EMB) genes required for growth and development in Arabidopsis. New Phytol. 226, 306–325. 10.1111/nph.16071

Michaels, S.D., Amasino, R.M., 1999. FLOWERING LOCUS C encodes a novel MADS domain protein that acts as a repressor of flowering. Plant Cell 11, 949–956. 10.1105/tpc.11.5.949

Millán-Zambrano, G., Burton, A., Bannister, A.J., Schneider, R., 2022. Histone post-translational modifications — cause and consequence of genome function. Nat. Rev. Genet. 23, 563–580. 10.1038/s41576-022-00468-7

Molitor, A.M., Bu, Z., Yu, Y., Shen, W.-H., 2014. Arabidopsis AL PHD-PRC1 Complexes Promote Seed Germination through H3K4me3-to-H3K27me3 Chromatin State Switch in Repression of Seed Developmental Genes. PLoS Genet. 10, e1004091. 10.1371/journal.pgen.1004091

Mouriz, A., López-González, L., Jarillo, J.A., Piñeiro, M., 2015. PHDs govern plant development. Plant Signal. Behav. 10, e993253. 10.4161/15592324.2014.993253

Ohad, N., Margossian, L., Hsu, Y.C., Williams, C., Repetti, P., Fischer, R.L., 1996. A mutation that allows endosperm development without fertilization. Proc. Natl. Acad. Sci. U. S. A. 93, 5319– 5324.

Ono, A., Kinoshita, T., 2021. Epigenetics and plant reproduction: Multiple steps for responsibly handling succession. Curr. Opin. Plant Biol., Epigenetics 61, 102032. 10.1016/j.pbi.2021.102032

Peng, L., Wang, L., Zhang, Y., Dong, A., Shen, W.-H., Huang, Y., 2018. Structural Analysis of the Arabidopsis AL2-PAL and PRC1 Complex Provides Mechanistic Insight into Active-to-Repressive Chromatin State Switch. J. Mol. Biol. 430, 4245–4259. 10.1016/j.jmb.2018.08.021

Pien, S., Fleury, D., Mylne, J.S., Crevillen, P., Inzé, D., Avramova, Z., Dean, C., Grossniklaus, U., 2008. ARABIDOPSIS TRITHORAX1 dynamically regulates FLOWERING LOCUS C activation via histone 3 lysine 4 trimethylation. Plant Cell 20, 580–588. 10.1105/tpc.108.058172

Quinlan, A.R., Hall, I.M., 2010. BEDTools: a flexible suite of utilities for comparing genomic features. Bioinforma. Oxf. Engl. 26, 841–842. 10.1093/bioinformatics/btq033

Rademacher, E.H., Lokerse, A.S., Schlereth, A., Llavata-Peris, C.I., Bayer, M., Kientz, M., Freire Rios, A., Borst, J.W., Lukowitz, W., Jürgens, G., Weijers, D., 2012. Different Auxin Response Machineries Control Distinct Cell Fates in the Early Plant Embryo. Dev. Cell 22, 211–222. 10.1016/j.devcel.2011.10.026

Ramírez, F., Dündar, F., Diehl, S., Grüning, B.A., Manke, T., 2014. deepTools: a flexible platform for exploring deep-sequencing data. Nucleic Acids Res. 42, W187–191. 10.1093/nar/gku365

Ratcliffe, O.J., Kumimoto, R.W., Wong, B.J., Riechmann, J.L., 2003. Analysis of the Arabidopsis MADS AFFECTING FLOWERING gene family: MAF2 prevents vernalization by short periods of cold. Plant Cell 15, 1159–1169. 10.1105/tpc.009506

Sablowski, R., 2007. Flowering and determinacy in Arabidopsis. J. Exp. Bot. 58, 899–907. 10.1093/jxb/erm002

Schmid, M.W., 2017. RNA-Seq Data Analysis Protocol: Combining In-House and Publicly Available Data. Methods Mol. Biol. Clifton NJ 1669, 309–335. 10.1007/978-1-4939-7286-9_24

Schuettengruber, B., Martinez, A.-M., Iovino, N., Cavalli, G., 2011. Trithorax group proteins: switching genes on and keeping them active. Nat. Rev. Mol. Cell Biol. 12, 799–814. 10.1038/nrm3230

Sheldon, C.C., Hills, M.J., Lister, C., Dean, C., Dennis, E.S., Peacock, W.J., 2008. Resetting of FLOWERING LOCUS C expression after epigenetic repression by vernalization. Proc. Natl. Acad. Sci. U. S. A. 105, 2214–2219. 10.1073/pnas.0711453105

Shu, J., Chen, C., Thapa, R.K., Bian, S., Nguyen, V., Yu, K., Yuan, Z.-C., Liu, J., Kohalmi, S.E., Li, C., Cui, Y., 2019. Genome-wide occupancy of histone H3K27 methyltransferases CURLY LEAF and SWINGER in *Arabidopsis* seedlings. Plant Direct 3, e00100. 10.1002/pld3.100

Simonini, S., Bemer, M., Bencivenga, S., Gagliardini, V., Pires, N.D., Desvoyes, B., van der Graaff, E., Gutierrez, C., Grossniklaus, U., 2021. The Polycomb group protein MEDEA controls cell proliferation and embryonic patterning in Arabidopsis. Dev. Cell 56, 1945–1960.e7. 10.1016/j.devcel.2021.06.004

Song, Y., Gao, J., Yang, F., Kua, C.-S., Liu, J., Cannon, C.H., 2013. Molecular Evolutionary Analysis of the Alfin-Like Protein Family in Arabidopsis lyrata, Arabidopsis thaliana, and Thellungiella halophila. PLOS ONE 8, e66838. 10.1371/journal.pone.0066838

Sridha, S., Wu, K., 2006. Identification of AtHD2C as a novel regulator of abscisic acid responses in Arabidopsis. Plant J. Cell Mol. Biol. 46, 124–133. 10.1111/j.1365-313X.2006.02678.x

Su, X.-M., Yuan, D.-Y., Liu, N., Zhang, Z.-C., Yang, M., Li, L., Chen, S., Zhou, Y., He, X.-J., 2025. ALFIN-like proteins link histone H3K4me3 to H2A ubiquitination and coordinate diverse chromatin modifications in Arabidopsis. Mol. Plant 18, 130–150. 10.1016/j.molp.2024.12.007

Sung, S., Schmitz, R.J., Amasino, R.M., 2006. A PHD finger protein involved in both the vernalization and photoperiod pathways in Arabidopsis. Genes Dev. 20, 3244–3248. 10.1101/gad.1493306

Tao, Z., Hu, H., Luo, X., Jia, B., Du, J., He, Y., 2019. Embryonic resetting of the parental vernalized state by two B3 domain transcription factors in Arabidopsis. Nat. Plants 5, 424–435. 10.1038/s41477-019-0402-3

ten Hove, C.A., Lu, K.-J., Weijers, D., 2015. Building a plant: cell fate specification in the early *Arabidopsis* embryo. Development 142, 420–430. 10.1242/dev.111500

Tian, L., Wang, J., Fong, M.P., Chen, M., Cao, H., Gelvin, S.B., Chen, Z.J., 2003. Genetic control of developmental changes induced by disruption of Arabidopsis histone deacetylase 1 (AtHD1) expression. Genetics 165, 399–409. 10.1093/genetics/165.1.399

Trachsel, C., Panse, C., Kockmann, T., Wolski, W.E., Grossmann, J., Schlapbach, R., 2018. rawDiag: An R Package Supporting Rational LC-MS Method Optimization for Bottom-up Proteomics. J. Proteome Res. 17, 2908–2914. 10.1021/acs.jproteome.8b00173

Van Bel, M., Silvestri, F., Weitz, E.M., Kreft, L., Botzki, A., Coppens, F., Vandepoele, K., 2022. PLAZA 5.0: extending the scope and power of comparative and functional genomics in plants. Nucleic Acids Res. 50, D1468–D1474. 10.1093/nar/gkab1024

Vielle-Calzada, J.P., Thomas, J., Spillane, C., Coluccio, A., Hoeppner, M.A., Grossniklaus, U., 1999. Maintenance of genomic imprinting at the Arabidopsis medea locus requires zygotic DDM1 activity. Genes Dev. 13, 2971–2982. 10.1101/gad.13.22.2971

Wang, Z.-P., Xing, H.-L., Dong, L., Zhang, H.-Y., Han, C.-Y., Wang, X.-C., Chen, Q.-J., 2015. Egg cell-specific promoter-controlled CRISPR/Cas9 efficiently generates homozygous mutants for multiple target genes in Arabidopsis in a single generation. Genome Biol. 16, 144. 10.1186/s13059-015-0715-0

Wasserzug-Pash, P., Klutstein, M., 2019. Epigenetic changes in mammalian gametes throughout their lifetime: the four seasons metaphor. Chromosoma 128, 423–441. 10.1007/s00412-019-00704-w

Wei, W., Zhang, Y.-Q., Tao, J.-J., Chen, H.-W., Li, Q.-T., Zhang, W.-K., Ma, B., Lin, Q., Zhang, J.-S., Chen, S.-Y., 2015. The Alfin-like homeodomain finger protein AL5 suppresses multiple negative factors to confer abiotic stress tolerance in Arabidopsis. Plant J. 81, 871–883. 10.1111/tpj.12773

Wu, K., Malik, K., Tian, L., Brown, D., Miki, B., 2000. Functional analysis of a RPD3 histone deacetylase homologue in Arabidopsis thaliana. Plant Mol. Biol. 44, 167–176. 10.1023/a:1006498413543

Xu, L., Wang, Y., Li, X., Hu, Q., Adamkova, V., Xu, J., Harris, C.J., Ausin, I., 2024. The H3K4me3 binding ALFIN-LIKE proteins recruit SWR1 for gene-body deposition of H2A.Z. 10.1101/2024.09.12.612642

Yoshida, N., Yanai, Y., Chen, L., Kato, Y., Hiratsuka, J., Miwa, T., Sung, Z.R., Takahashi, S., 2001. EMBRYONIC FLOWER2, a novel polycomb group protein homolog, mediates shoot development and flowering in Arabidopsis. Plant Cell 13, 2471–2481. 10.1105/tpc.010227

Yu, F., Haynes, S.E., Nesvizhskii, A.I., 2021. IonQuant Enables Accurate and Sensitive Label-Free Quantification With FDR-Controlled Match-Between-Runs. Mol. Cell. Proteomics MCP 20, 100077. 10.1016/j.mcpro.2021.100077

Yun, M., Wu, J., Workman, J.L., Li, B., 2011. Readers of histone modifications. Cell Res. 21, 564–578. 10.1038/cr.2011.42

Zenk, F., Loeser, E., Schiavo, R., Kilpert, F., Bogdanović, O., Iovino, N., 2017. Germ line-inherited H3K27me3 restricts enhancer function during maternal-to-zygotic transition. Science 357, 212–216. 10.1126/science.aam5339

Zhang, Y., Liu, T., Meyer, C.A., Eeckhoute, J., Johnson, D.S., Bernstein, B.E., Nusbaum, C., Myers, R.M., Brown, M., Li, W., Liu, X.S., 2008. Model-based analysis of ChIP-Seq (MACS). Genome Biol. 9, R137. 10.1186/gb-2008-9-9-r137

Zhou, Y., Wang, Y., Krause, K., Yang, T., Dongus, J.A., Zhang, Y., Turck, F., 2018. Telobox motifs recruit CLF/SWN–PRC2 for H3K27me3 deposition via TRB factors in Arabidopsis. Nat. Genet. 50, 638–644. 10.1038/s41588-018-0109-9

